# Treatment of pancreatic cancer with irreversible electroporation and intratumoral CD40 antibody stimulates systemic immune responses that inhibit liver metastasis in an orthotopic model

**DOI:** 10.1101/2022.10.04.510847

**Authors:** Jayanth S. Shankara Narayanan, Tomoko Hayashi, Suna Erdem, Sara McArdle, Herve Tiriac, Partha Ray, Minya Pu, Zbigniew Mikulski, Aaron Miller, Karen Messer, Dennis Carson, Stephen P. Schoenberger, Rebekah R. White

## Abstract

**Background:** Pancreatic cancer (PC) has a poor prognosis, and most patients present with either locally advanced or distant metastatic disease. Irreversible Electroporation (IRE) is a non-thermal method of ablation used clinically in locally advanced PC, but most patients eventually develop distant recurrence. We have previously shown that IRE alone is capable of generating protective, neoantigen-specific immunity. Here we aim to generate meaningful therapeutic immune effects by combining IRE with local (intratumoral) delivery of a CD40 agonistic antibody (CD40Ab).

**Methods:** KPC46 organoids were generated from a tumor-bearing male KrasLSL-G12D-p53LSL-R172H-Pdx-1-Cre (KPC) mouse. Orthotopic tumors were established in the pancreatic tail of B6/129 F1J mice via laparotomy. Mice were randomized to treatment with either sham laparotomy, IRE alone, CD40Ab alone, or IRE followed immediately by CD40Ab injection. Metastatic disease and immune infiltration in the liver were analyzed 14 days post-procedure using flow cytometry and multiplex immunofluorescence imaging with spatial analysis. Candidate neoantigens were identified by mutanome profiling of tumor tissue for *ex vivo* functional analyses.

**Results:** The combination of IRE+CD40Ab improved median survival to greater than 35 days, significantly longer than IRE (21 days) or CD40Ab (24 days) alone (p<0.01). CD40Ab decreased metastatic disease burden, with less disease in the combination group than in the sham group or IRE alone. Immunohistochemistry of liver metastases revealed a more than two-fold higher infiltration of CD8+ T-cells in the IRE+CD40Ab group than in any other group (p<0.01). Multiplex immunofluorescence imaging revealed a 4-6-fold increase in the density of CD80+CD11c+ activated dendritic cells (p<0.05), which were spatially distributed throughout the tumor unlike the sham group, where they were restricted to the periphery. In contrast, CD4+FoxP3+ T-regulatory cells (p<0.05) and Ly6G+ MDSCs (P<0.01) were reduced and restricted to the tumor periphery in the IRE+CD40Ab group. T-cells from the IRE+CD40Ab group recognized significantly more peptides representing candidate neoantigens than did T-cells from the IRE or untreated control groups.

**Conclusions:** IRE can induce local tumor regression and neoantigen-specific immune responses. Addition of CD40Ab to IRE improved dendritic cell activation and neoantigen recognition, while generating a strong systemic anti-tumor T-cell response that inhibited metastatic disease progression.

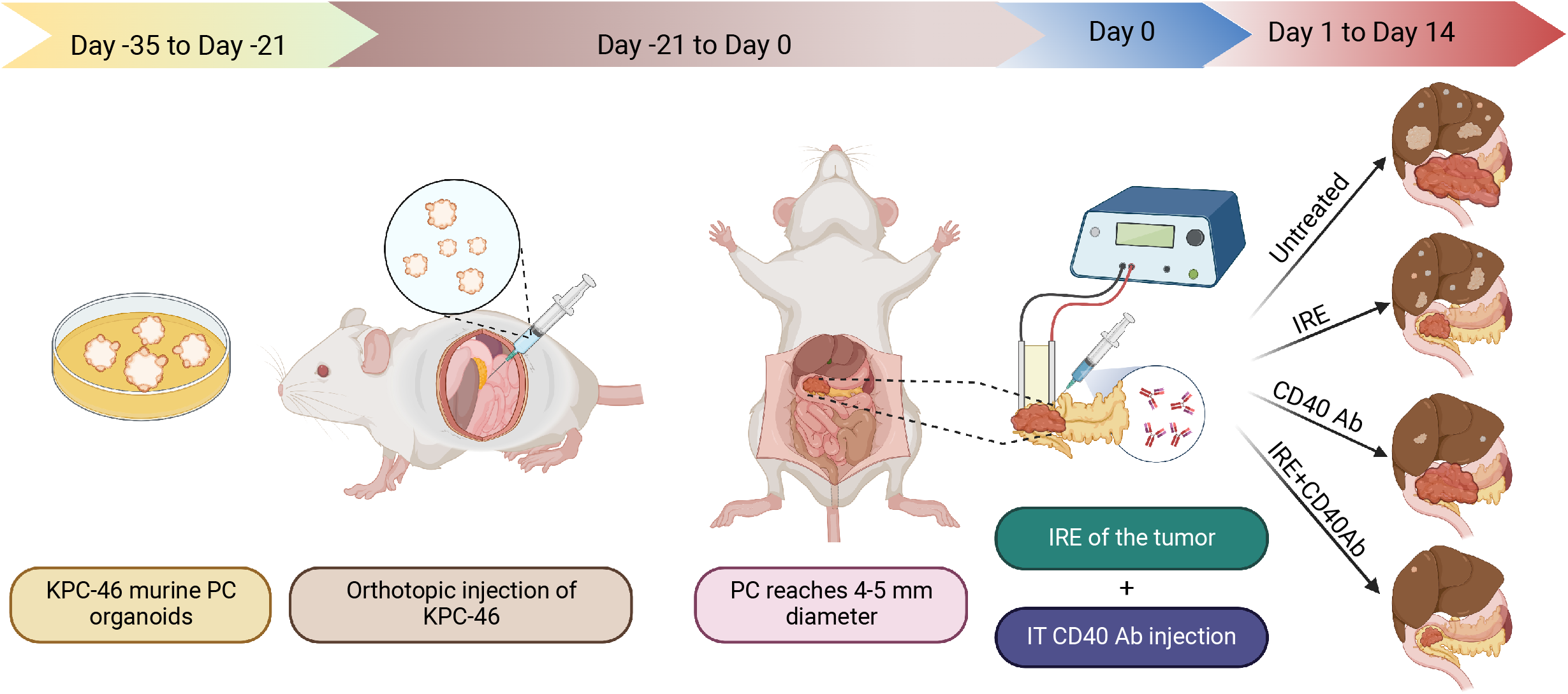

## INTRODUCTION

Approximately one third of the over 50,000 people diagnosed with pancreatic cancer (PC) in the U.S. each year will have locally advanced disease that appears localized radiographically but is not amenable to complete surgical resection [1]. The majority of these patients have occult micrometastatic disease [2], and there is nearly uniform agreement that patients with good performance status should receive systemic chemotherapy as their first line of therapy [3]. Although many of these patients will progress on chemotherapy, up to half initially attain disease control on multi-agent chemotherapy regimens [4, 5]. The principal challenge for these patients is that disease control is not sustainable, either due to acquired resistance or cumulative toxicity. Local therapies are appealing for such patients with localized but unresectable disease.

Irreversible electroporation (IRE) is a non-thermal method of inducing tumor cell death without destruction of adjacent vascular structures [6]. Unlike thermal ablation techniques, such as radiofrequency ablation (RFA), IRE is not vulnerable to “heat sink” effects, in which blood flowing through adjacent blood vessels decreases the effectiveness of ablation [7]. Specific to PC, the acellular structural components of blood vessels are not damaged by IRE [8], which permits safe treatment of tumors with vascular involvement. Multiple individual institutions [9-11] and a few multicenter studies [12-15] have published clinical experiences with IRE for locally advanced PC. These nonrandomized studies suggest that IRE is associated with longer survival than standard therapy alone. However, the fundamental problem with any local therapy for PC is that most patients develop distant recurrence, which highlights the need for better adjunctive treatment of micrometastatic disease [12, 15-17].

Our overarching hypothesis is that tumor ablation induces anti-tumor immune responses by increasing the availability of tumor-specific neoantigens (NeoAg’s) in an inflammatory context. NeoAg’s released by the tumor are processed by local antigen presenting cells (APCs) and stimulate adaptive immune responses. Several studies—including our own—have specifically examined immune responses to IRE [18-26]. Theoretically, IRE may be more effective at generating immune responses than thermal ablation due to greater preservation of protein antigens and the vascular structures that allow immune cell infiltration [27]. We have previously shown that IRE alone is effective at achieving local tumor control and protective immunity to tumor rechallenge in an immunocompetent mouse model, in which a cell line established from a genetically engineered “KPC” mouse [28] was used to generate subcutaneous (SQ) tumors [29].

Similar to prior studies in other tumor types, we found that IRE requires an intact immune system to be effective [18, 20, 24]. Depletion and adoptive transfer experiments identified CD8+ and CD4+ T-cells as the most important effectors of anti-tumor response after IRE [24]. CD8^+^ T cells from IRE-responsive mice were reactive against peptides representing model-inherent “NeoAg’s” that were identified by next generation sequencing (NGS) of nucleic acids derived from the tumor [24]. In this SQ model, the triple combination of IRE with intratumoral toll-like receptor-7 (TLR7) agonist (1V270) and systemic anti-programmed death-1 receptor (PD)-1 checkpoint inhibition resulted in improved local tumor control and also resulted in elimination of small, untreated tumors on the contralateral flank. Neither the addition to IRE of systemic anti-PD1 Ab alone nor local TLR7 agonism alone produced these effects [24]. These data demonstrate proof-of-principle that a combination of IRE with agents that enhance the innate and adaptive immune systems can potentially produce therapeutic immune responses (i.e., “abscopal” effects).

The CD40 receptor is expressed predominantly on antigen-presenting cells (APCs). Its activation co-stimulates antigen-loaded APCs, which promotes maturation and antigen presentation, acting as a bridge between the innate and adaptive immune systems [30, 31]. As such, CD40 agonists potentially can synergize with checkpoint inhibitors, which act on adaptive immune cells. Similar to the disappointing results with checkpoint inhibitors in humans with PC [32, 33], checkpoint inhibitors alone have not been effective in mouse models of PC [21, 34]. However, CD40 agonistic antibodies (CD40Ab) were able to help overcome resistance to checkpoint inhibition in KPC mice [34], and several CD40 agonists are being studied clinically in combination with chemotherapy and/or checkpoint inhibition [35, 36]. There are potential advantages to local delivery of CD40 agonists directly to the source of the NeoAg’s (the tumor) while potentially limiting systemic adverse side effects. We hypothesized that local delivery of CD40Ab at the time of IRE would improve immune responses to IRE. We used orthotopic mouse models of pancreatic cancer that recapitulate the human disease to demonstrate that this “all local” treatment combination not only improved local tumor control but also generated therapeutic immune responses that decreased metastatic disease burden.

## RESULTS

### Local delivery of agonistic CD40Ab following IRE improves local tumor control and generates abscopal effects in a SQ model of PC

To screen for activity as adjuvant immunotherapy, we treated SQ KPC4580P tumors by IRE in combination with CD40 Ab. The combination of IRE and CD40Ab resulting in 6 of 10 complete responders and 3 more showing only limited tumor progression (Fig. 1A). Similar to our prior studies, IRE alone induced complete responses in only 2/10 mice in this model. CD40Ab alone did not induce any complete responses. By comparison, several other immunomodulatory agents such as anti-PD-1 TLR7, TLR9 and STING agonists were tested on the same model with no significant effects as single agents (Supplementary Fig. 1).

**Figure 1.**
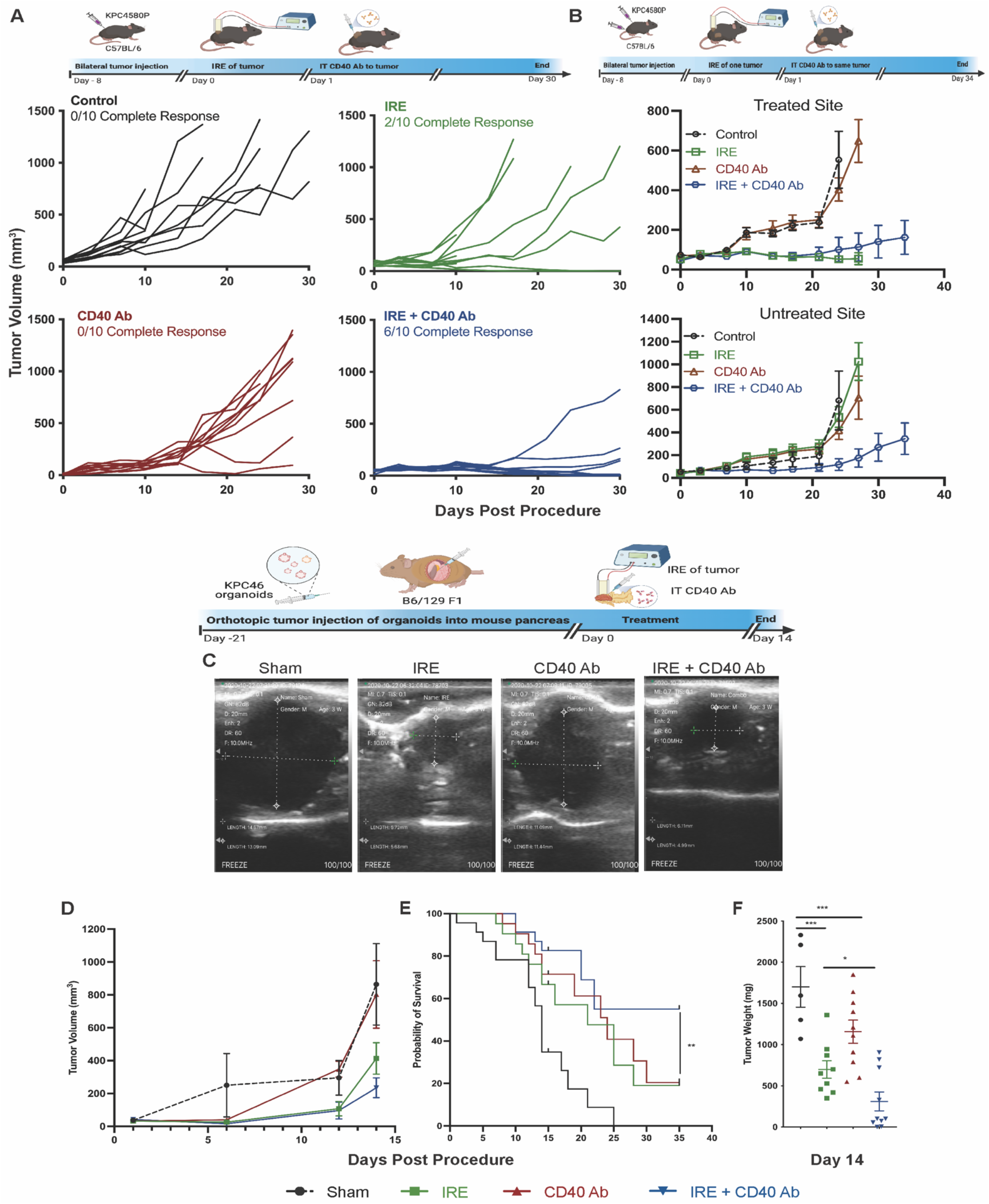
Combination of IRE with agonistic CD40Ab reduces tumor growth and improves survival in subcutaneous and orthotopic organoid PDAC mouse model. A – Subcutaneous KPC4580P tumor growth in C57BL/6 (n=10/group) mice implanted on one side of the flanks at 8 days before IRE (5 × 10^5^ cells/mouse). Each data point represents tumor volume of a single mouse followed along the growth curve. B - Tumor growth curves of bilateral subcutaneous KPC4580P tumors at the treated site (top) and untreated site (bottom), of n = 5 mice/group, implanted 8 days before IRE, followed by IT CD40 Ab injection on Day 1. Tumor volumes were measured using calipers and plotted as mean ± SEM. C – 30,000 murine PDAC organoids KPC46 were injected into the pancreas of B6/129 F1 hybrid mice 21 days before treatment. Ultrasound monitoring of orthotopic organoid tumors Day 7 post treatment showing responses to treatment. Dotted lines show the dimensions of the orthotopic tumors under sltrasound imaging. D –-Tumor volumes were measured using ultrasound and plotted as mean ± SEM of n = 10 mice/group. E – Kaplan meier survival analysis of orthotopic organoid KPC46 tumor bearing mice post treatment (n=20 mice/group) cumulative of 3 independent experiments showing significant survival benefit offered by IRE+CD40Ab combination **, p<0.01 by log rank test. F – Tumor weights as a measure of primary tumor burden, tumors were excised from mice upon euthanasia, 14 days post treatment, each data point represents single tumor weight represented as mean ± SEM. *, P < 0.05; **, P < 0.01; ***, P < 0.001 by one-way ANOVA with post hoc Benferroni test.

To assess the systemic effects of this combination, we utilized a model harboring small bilateral, symmetric flank tumors. When one side was treated with IRE and/or CD40 Ab injection, both IRE and IRE+CD40Ab were equally effective in controlling the local (treated) tumor (Fig. 1B). IRE alone had no effect on growth of the untreated tumor on the contralateral flank. However, when IRE was followed by local delivery of CD40Ab, not only did the treated tumor shrink, but the growth of untreated tumors on the contralateral flank was also significantly inhibited (p <0.01, Fig. 1B). Further, 3 of 9 animals in this group showed complete tumor regression on both the treated as well as the untreated site (Fig. 1B). In order to assess the prophylactic function of this anti-tumor immune response, we rechallenged the complete responders from the single tumor model with live tumor cells 20 days post-procedure. None of the complete responders in either the IRE alone or the IRE+CD40Ab group was able to support the growth of the secondary tumor, demonstrating an active systemic anti-tumor immune rejection. We also performed a delayed rechallenge on these complete responders 200 days post-procedure and observed similar tumor rejection (Supplementary Fig. 2).

**Figure 2.**
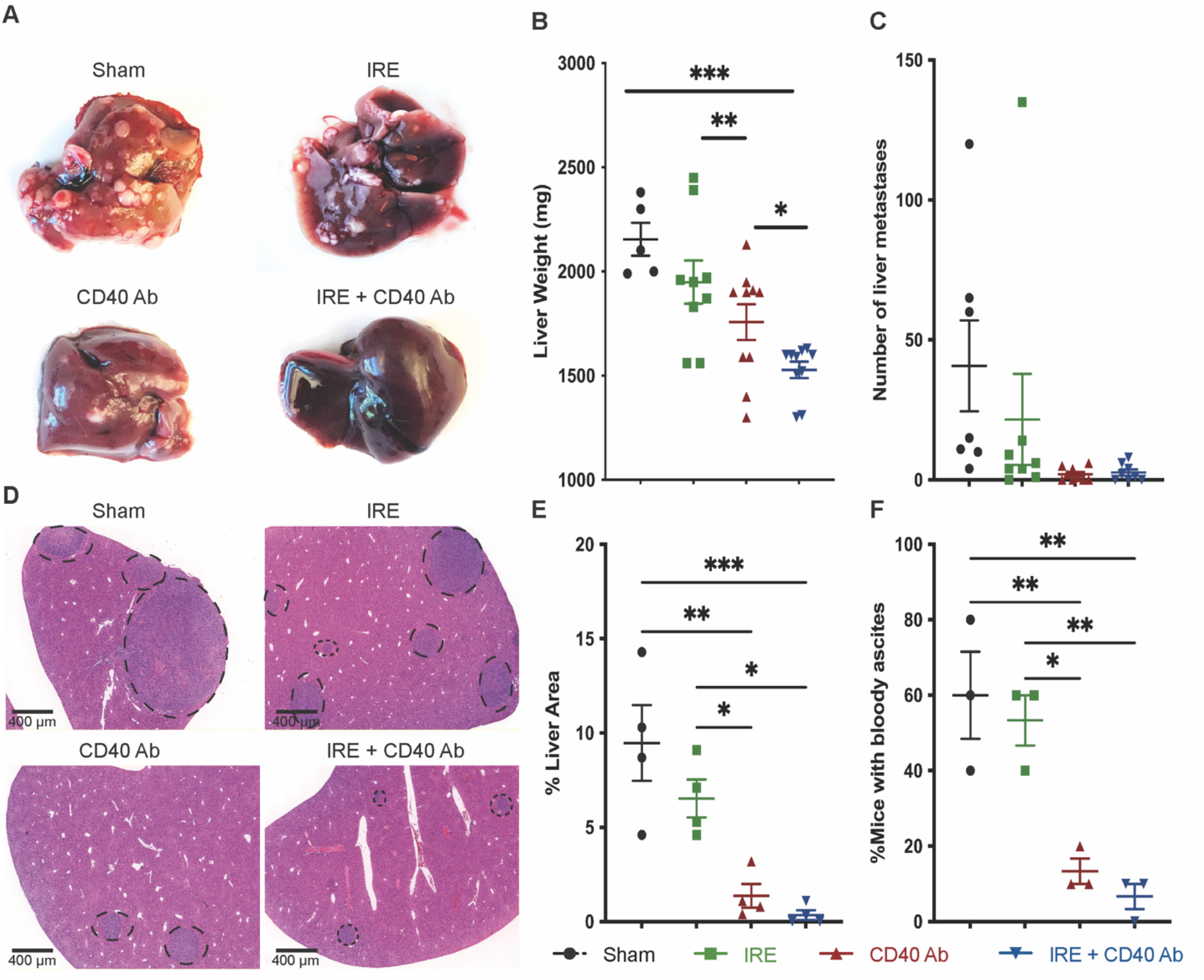
Combining IRE with CD40 agonistic activity inhibits tumor metastasis to the liver. A – Images of liver showing differences in metastasis. Whole livers were excised from orthotopic organoid KPC46 tumor bearing B6129 F1 mice upon euthanasia 14 days post treatment under different groups (n=10/group). B - Liver weights as a measure of metastatic burden with a significant reduction in metastasis seen with IRE+CD40Ab combination, weights of whole liver was measured in mice surviving 14 days post procedure each data point represents single liver weight, represented as mean ± SEM, E - Manual counting of visible metastatic nodules on the liver as a measure of metastatic tumor burden D and E histological examination of mice liver showing metastatic tumor nodules observed using H&E. Images were acquired using Zeiss slide scanner at 20X objective and the % liver area occupied by the tumor was calculated using QuPath plotted as mean ± SEM 3 sections/mouse, 4 mice/group. F - Untreated mice are more likely to develop bloody ascites upon tumor progression compared to CD40Ab or IRE+CD40Ab treatment. Each dot represents percentage of mice with bloody ascites/experimental group plotted as a mean of 3 independent experimental rounds. *, P < 0.05; **, P < 0.01; ***, P < 0.001; ns, not significant, by one-way ANOVA with post hoc Benferroni multiple comparison test.

### Local delivery of agonistic CD40Ab following IRE improves local tumor control and survival in an orthotopic organoid model of pancreatic cancer

The KPC-46 organoid cell line was derived from an aggressive tumor in a KPC mouse. When injected orthotopically into the pancreas, mice consistently develop primary tumors and liver metastases within a few weeks, modeling human PC. Since the liver is the most important site of metastatic disease in PC, we evaluated IRE+CD40Ab in this model. Serial ultrasound estimates of tumor volumes weekly post-procedure (Fig. 1C) suggest growth inhibition in tumors treated with IRE and IRE+CD40Ab (Fig. 1D). The lack of significant difference in the tumor volumes between the groups was possibly due to the inability of ultrasound to distinguish between viable tumor and scar tissue in the orthotopic setting. We observed similar effects in an orthotopic transplant model using the KPC4580P cell line [24] (Supplementary Fig. 3). However, when mice were sacrificed on Day 14 post-procedure, mean tumor weights were significantly (P<0.01) lower in the IRE (698 ± 106 mg) and IRE+CD40Ab (311 ± 114 mg) groups than in the sham (1700 ± 247 mg) or CD40 Ab alone (1160 ± 141 mg) groups (Fig. 1 F). Tumor weights were also significantly lower with IRE+CD40Ab than with IRE alone (p<0.05). In a separate survival experiment, sham-treated mice exhibited a median survival of only 14 days post-procedure (35 days after tumor implantation) due to rapid development of metastasis and increasing tumor burden. IRE alone or CD40 Ab alone improved median survival to 21 and 24 days, respectively, but these were significantly (p<0.01) shorter than the median survival of >35 days achieved by IRE+CD40Ab (Fig. 1E). Further, tumor immune-profiling revealed a trend toward increased infiltration of CD8+ T-cells into the primary tumors in all treatment groups compared to sham, which were predominantly positive for IFNγ (>40%) suggestive of robust anti-tumor cytotoxic activity (Supplementary Fig. 4).

### IRE in combination with CD40Ab inhibits metastatic tumor progression in the liver

We assessed metastatic disease burden in mice with orthotopic KPC-46 organoid tumors following treatment in multiple ways. Fig. 2A depicts the differences in the gross appearance of representative livers from each treatment group. Mean liver weights in the IRE+CD40Ab group were significantly lower than the sham (p<0.01) or IRE alone (p<0.05) groups (Fig. 2B). Manual counting of visible metastatic nodules did not reveal a significant difference in the number of macro metastases in mice treated with IRE+CD40Ab (Fig. 2C). However, histopathological evaluation of liver tissue (Fig. 2D) suggests a decrease in the size of liver metastases with the combination treatment. Quantification of the percentage of liver cross-sectional area occupied by tumor revealed a significant decrease in mice treated with IRE+CD40Ab or CD40Ab alone (Fig. 3E). Similarly, mice treated with IRE+CD40Ab or CD40Ab alone were less likely to have bloody peritoneal ascites than sham or IRE-treated mice (∼15% vs 60% over 3 experiments, Fig. 3F) as a secondary marker for metastatic progression. IRE alone had no effect on distant progression, consistent with our observations in the bilateral flank tumor model (Fig. 1B). We directly compared route of CD40Ab delivery in mice that were not treated with IRE and observed that IT injections of CD40Ab resulted in significantly better control of local tumor burden (p < 0.05) than systemic (intraperitoneal) delivery of CD40Ab without significantly affecting its ability to control intraperitoneal spread (Supplementary Fig. 5A&B). Interestingly, delayed (2 days post-IRE) systemic delivery of CD40Ab did not improve outcomes and was associated with increased metastatic progression in the liver compared to IT injection on the day of IRE (Supplementary Fig. 6C,D &E). These findings demonstrate that IRE alone inhibited local tumor growth, and local delivery of CD40 Ab alone inhibited metastatic progression. The combination had at least additive effects, effectively inhibiting both local and metastatic disease.

**Figure 3.**
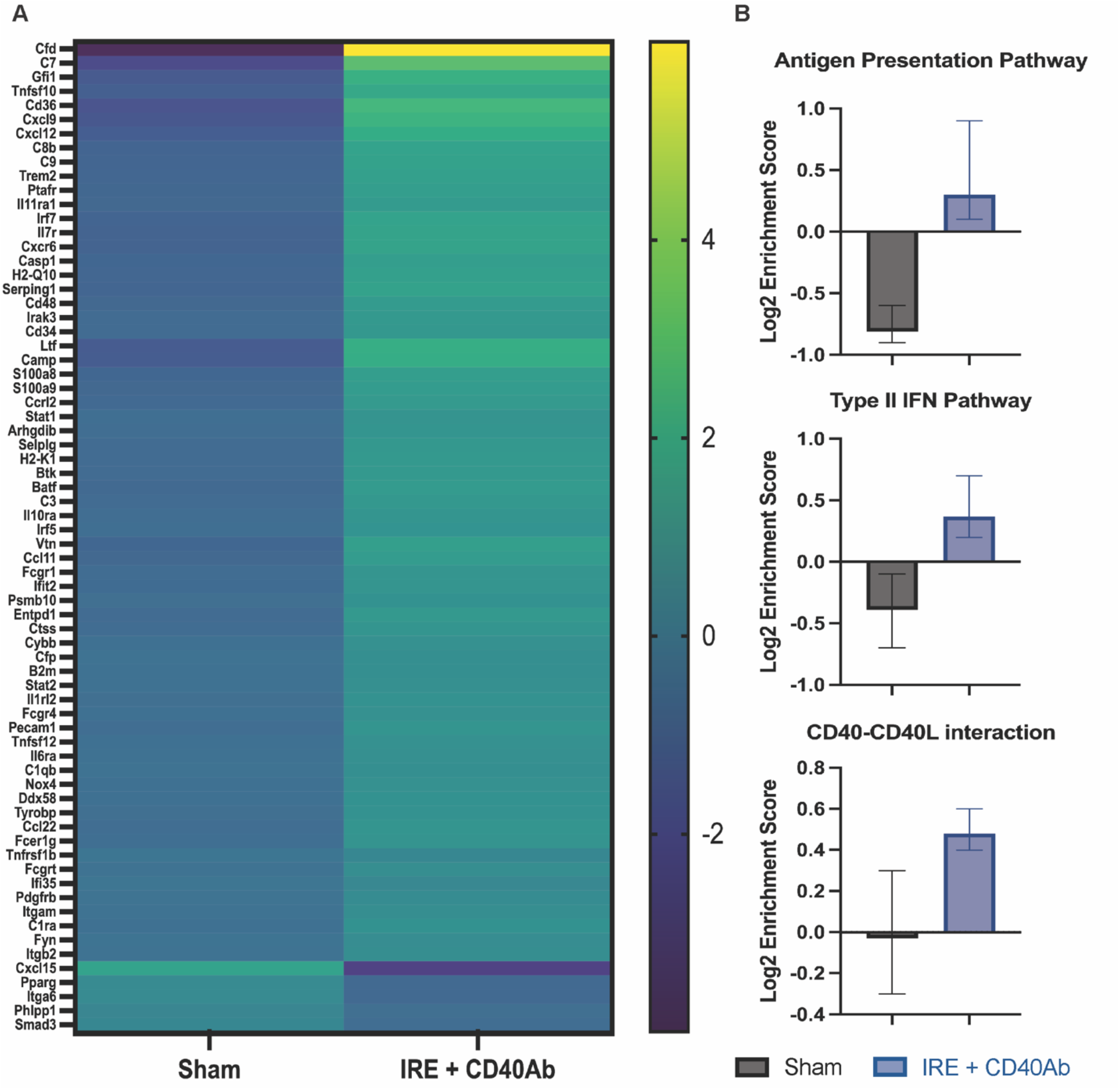
Activation of anti-tumor immune signaling pathways observed both locally and systemically upon treatment with combination of IRE and CD40Ab. A – RNA-Seq analysis of important immunoregulatory genes in the primary pancreatic tumor between the 2 groups. Heat map represents fold change in the expression levels of critical genes within tumor microenvironment between sham and IRE+CD40Ab groups (n=3/group) B – Gene set enrichment analysis of immunological pathway related genes, represented as Log2 enrichment scores of antigen presentation pathway (Top), IFN signaling pathway (Middle) and CD40 – CD40 L interaction pathway (Right) as mean ± SD of n=3/group. Statistical analysis was performed using 2-tailed student t-test. *, P < 0.05; **, P < 0.01; ***, P < 0.001.

### Gene expression changes following treatment with IRE+CD40Ab indicate an enhanced activation of antigen processing machinery

To assess whether treatment with IRE+CD40 Ab resulted in changes in the adaptive immune response within the primary tumor, bulk RNA-seq followed by gene set enrichment pathway analyses (GSEA) were performed. Fig. 3A shows a heatmap of gene expression changes that were significant post-treatment among selected immunomodulatory genes. In addition to the elevation of multiple complement pathway-related genes, several genes responsible for lymphocyte activation such as Irf7, Cd48, Stat1 and Stat2 were upregulated. Genes responsible for antigen presentation, such as Batf, which promotes antigen cross-presentation were also upregulated upon treatment with IRE+CD40Ab. Genes with immunosuppressive potential, such as Cxcl15, Pparg and Smad3, were significantly downregulated. The results of GSEA and gene ontology also indicated that several adaptive immune pathways were significantly enriched in the treatment group, such as the antigen presentation pathway and type-1 interferon pathway (Fig. 3B). As an indication of effective CD40 activation, overall downstream signalling resulting from the CD40-CD40L pathway was enriched in the tumor (Fig. 3B).

### Local treatment of pancreatic tumors with IRE+CD40Ab modulates the immune microenvironment within the liver

Reduction in metastatic disease could simply be the result of improved local tumor control and decreased metastatic spread to the liver. To determine whether the reduction in metastatic disease was immune-mediated, we examined immune infiltrates both in metastatic lesions and in bulk liver. Analysis of pan-immune cell marker CD45 by IHC (Fig. 4A, B) shows IRE+CD40Ab treatment resulted in an at least 4-fold higher density of CD45+ cells than IRE alone (1020 ± 192 vs 181 ± 13 cells/mm^2^, P<0.001) and 2-fold higher than CD40Ab alone (478 ± 31 cells/mm^2^, P<0.01). Further, infiltrating immune cells in tumors treated with IRE+CD40 Ab had a higher percentage of effector cytotoxic CD8+ T-cells (Fig. 4 A, C) than either CD40Ab alone (p <0.05), IRE (p <0.001), or sham treatment (p < 0.0001). Flow cytometry analysis of bulk liver tissue also showed an interesting increase in CD4+ T cell populations in the liver of treated mice (Fig. 4D) with corresponding decreases in the immune-suppressive myeloid derived suppressor cell (MDSC) population and T–regulatory cell (T-regs) population. Bulk liver flow cytometry did not reflect the changes seen in the CD8+ cytotoxic T-cell populations within tumors by IHC (Fig. 4A), with the images indicating that the changes in immune cell populations within the liver are restricted to metastatic tumor sites with minimal observed changes in the surrounding normal tissue. Further flow cytometry analysis revealed an increase in cross-presenting CD8^+^CD11c^+^ dendritic cells in the bulk liver (Fig. 4E).

**Figure 4.**
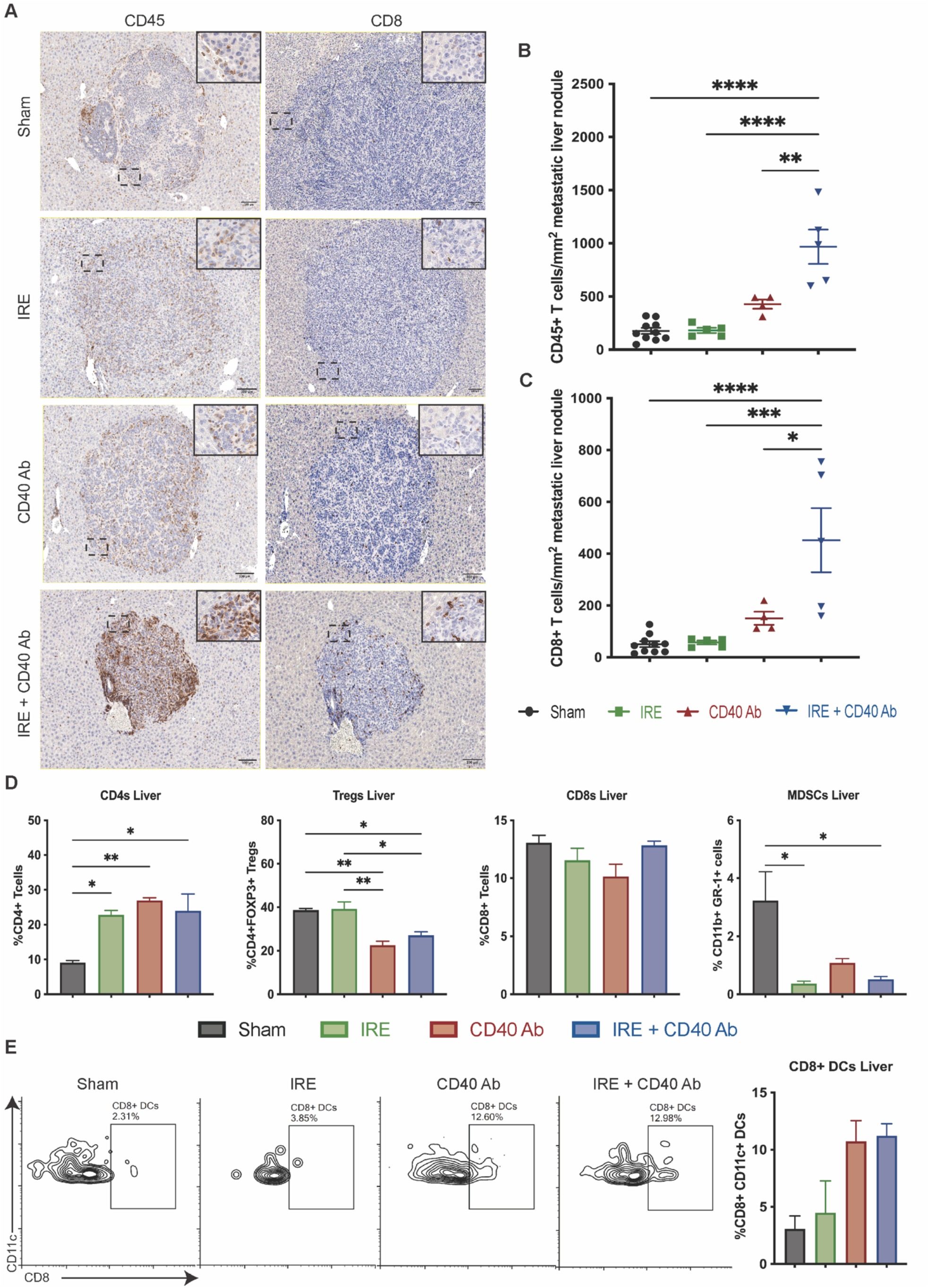
Immune infiltration in the liver shows that the combination of IRE and CD40 Ab induces an immune mediated prevention of metastatic spread. A - IHC for CD45 and CD8 in the liver showing increased specific staining within the metastatic tumor nodule, Images obtained using Leica Aperio slide scanner with 20X objective. Scale bar: 100 μm. B - Quantification of CD45 and CD8 positive cells respectively, within the metastatic tumor nodules of the liver was performed using QuPath 3.0, 3 mice/group including atleast 5 tumor nodules per group plotted as mean ± SEM. C – Flow cytometry of immune infiltrates in the metastatic site (bulk liver) performed on day 14 on whole liver (n=3/group) Decreased MDSCs and Tregs with the increased infiltration of CD4+ T cells cells suggest an anti-tumor immune activity at the distant metastatic site upon treatment with IRE+CD40Ab. D – Flow cytometry analysis of liver showing increased presence of cross-presenting CD8+ dendritic cells. Graphs plotted as mean ± SEM. *, P < 0.05; **, P < 0.01; ***, P < 0.001; by one-way ANOVA with post hoc Benferroni multiple comparison test. Gating strategy is described in Supplementary data.

To further assess the immunological changes taking place at the distant metastatic site, we performed multiplex immunofluorescence analysis followed by fluorescence intensity quantification for each channel on a “per cell” basis. We observed that tumor deposits, identified by dense pan-cytokeratin (PanCK, green – Fig. 5A) signal, were less frequent and smaller in size in liver sections from mice treated with CD40Ab alone or IRE+CD40Ab, consistent with our previous analysis with H&E stains. Further, we observed that the presence of CD11c+ dendritic cells (red) within tumors increased following treatment with CD40Ab alone or IRE+CD40Ab. There was a concomitant increase in CD80+ (Cyan) CD11c+ activated dendritic cells. Quantification (Fig. 5B, C) revealed a 4-6 fold increase in the density of CD11c as well as CD80+CD11c+ activated dendritic cells following IRE+CD40Ab treatment which was statistically significant (p < 0.05). This increase in activated CD80^+^CD11c^+^dendritic cells, which are professional antigen presenting cells, supports our earlier observation of increased expression of antigen-presentation pathway genes and CD40 – CD40 L interaction pathway genes from RNAseq analysis of primary tumors (Fig. 3B). On serial sections containing the same metastatic deposits we also observed a decrease in CD4+ FoxP3+ (red and cyan, Fig. 5D, E) T-regulatory cells (p< 0.05) whereas total CD4+ cells increased (not shown), resulting in an increased T-helper to T-regulatory cell ratio (CD4 : T-reg ratio, Fig. 5G) towards a less immunosuppressive niche. Ly6G+ (yellow) immunosuppressive MDSCs were not significantly reduced in number (Fig. 5D,F). There was a statistically significant increase (P <0.05) in F4/80-positive macrophages (green) within metastatic deposits (Fig. 5H) with flow cytometry data indicating that these macrophages are polarized towards the M1 phenotype (Supplementary Fig. 6). Together, these data indicate that local treatment of the primary tumor in the pancreas with CD40 Ab increases the density and activity of antigen-presenting cells within distant liver metastases.

**Figure 5.**
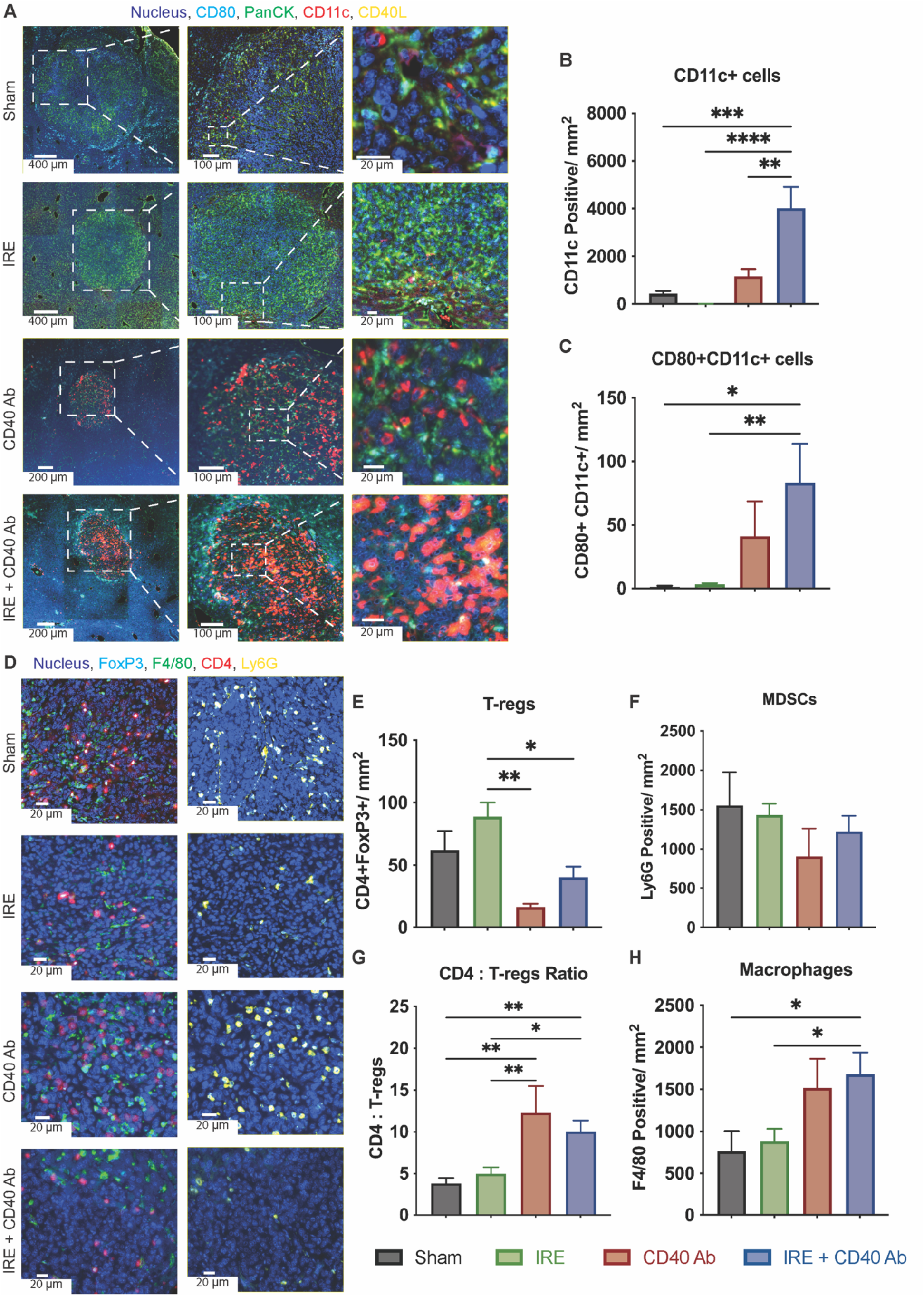
Multiplex immunofluorescence imaging shows an anti-tumor immune microenvironment within metastatic liver nodules post treatment with IRE+CD40Ab. A – Representative multiplex fluorescence microscopy images on FFPE tissue sections of mice liver showing metastatic nodes showing tumor infiltrating immune cells stained with either Panel 1 – CD11c:Alexa 647(red), CD80:Alexa 488(Cyan), CD40 L:Alexa 790 (Yellow), PanCK:Alexa 555(Green), DAPI (Blue), Scale bars set at 400 om (Low magnification Sham and IRE), 200 om (Low magnification CD40 Ab and IRE+CD40Ab), 100 om (Medium magnification) and 20 om (High magnification). B and C - Quantification of dendritic cell (CD11c+) and activated dendritic cell (CD80+CD11c+) infiltration per mm2 of the metastatic nodules was performed using QuPath 3.0 software using atleast 5 different nodules spanning 3 biological replicates per group. D-Representative multiplex immunofluorescence imaging of CD4 Alexa 647) (Red), F4/80 – Alexa 555 (Green), Ly6G – Alexa 790 (Yellow), FoxP3 – 488 (Cyan), DAPI (Blue). Scale bars set at 20 om). E - H – Quantification of T regulatory cell (CD4+FoxP3+), Macrophage (F4/80+) and Myeloid Derived Suppressor Cell (MDSC – Ly6G+) infiltration to the metastatic nodules was performed using QuPath 3.0 software using atleast 5 different nodules spanning 3 biological replicates per group. The number of immune cells was normalized to the area of the nodes and presented as mean ± SEM. *, P < 0.05; **, P < 0.01; ***, P < 0.001; by one-way ANOVA with post hoc Benferroni multiple comparison test.

### IRE+CD40Ab combination therapy generates a broader recognition of tumor NeoAg’s by T-cells

In our previous study using the KPC4580P cell line, we had shown that IRE stimulates T-cell responses against tumor-specific alloantigens (model NeoAg’s) [24]. Following the observation of increased CD8+ T-cell infiltration within metastatic tumor sites, we explored whether the combination of IRE and CD40Ab can induce similar or stronger responses than IRE alone in a different less immunogenic, orthotopic mouse model. RNA and DNA were isolated from KPC-46 organoid orthotopic tumors, and expressed non-synonymous mutations (against wild-type B6/129 F1 hybrid background) were identified using whole exome sequencing and RNAseq. Identified variants are depicted in a Circos plot (Fig. 6A). A total of 58 variants were prioritized based on their high RNA expression and sequencing depth, and 116 peptides were tested in 10 pools of 8-12 peptides each for their ability to induce IFNγ secretion in ELISPOT assay. Fig. 6B shows a representative ELISPOT round where we observed that T-cells from mice treated with IRE alone had significantly increased reactivity over background response against non-specific peptides, against pools 2, 7 and 9. T-cells from the IRE+CD40Ab group showed significantly increased reactivity against pools 1, 2 and 3. Untreated tumor-bearing mice did not show significant reactivity to any of the tested pools. Although the trend was similar between mice and between repeated experiments, a consistent significant hit on a single peptide pool was not achieved with either of the treatment groups. Deconvolution was performed on peptide pool 2 alone, which was recognized by 60% mice in the IRE+CD40Ab group per experiment on average and by 40% mice in the the IRE alone group over 3 experiments. In a representative ELISPOT experiment on pool 2 (Fig. 6C), we observed that T cells isolated from mice treated with IRE+CD40Ab recognized more peptides (7/12) than the those mice treated with IRE alone (2/12) or control mice (1/12). This trend was consistent across experiments, where T-cells from mice treated with IRE+CD40Ab positively recognized more (65 ± 9.3%) candidate neoantigens per round than IRE alone (36.9 ± 6.2%) or even the mice vaccinated with lethally irradiated KPC-46 organoids (Fig. 6D). The intensities of positive recognitions were higher with the combination but not statistically different between the treatment groups (data not shown). These data show that the addition of CD40Ab preserved the “in-situ” anti-tumor vaccination effect induced by IRE and also increased the breadth of NeoAg recognition generated by IRE.

**Figure 6.**
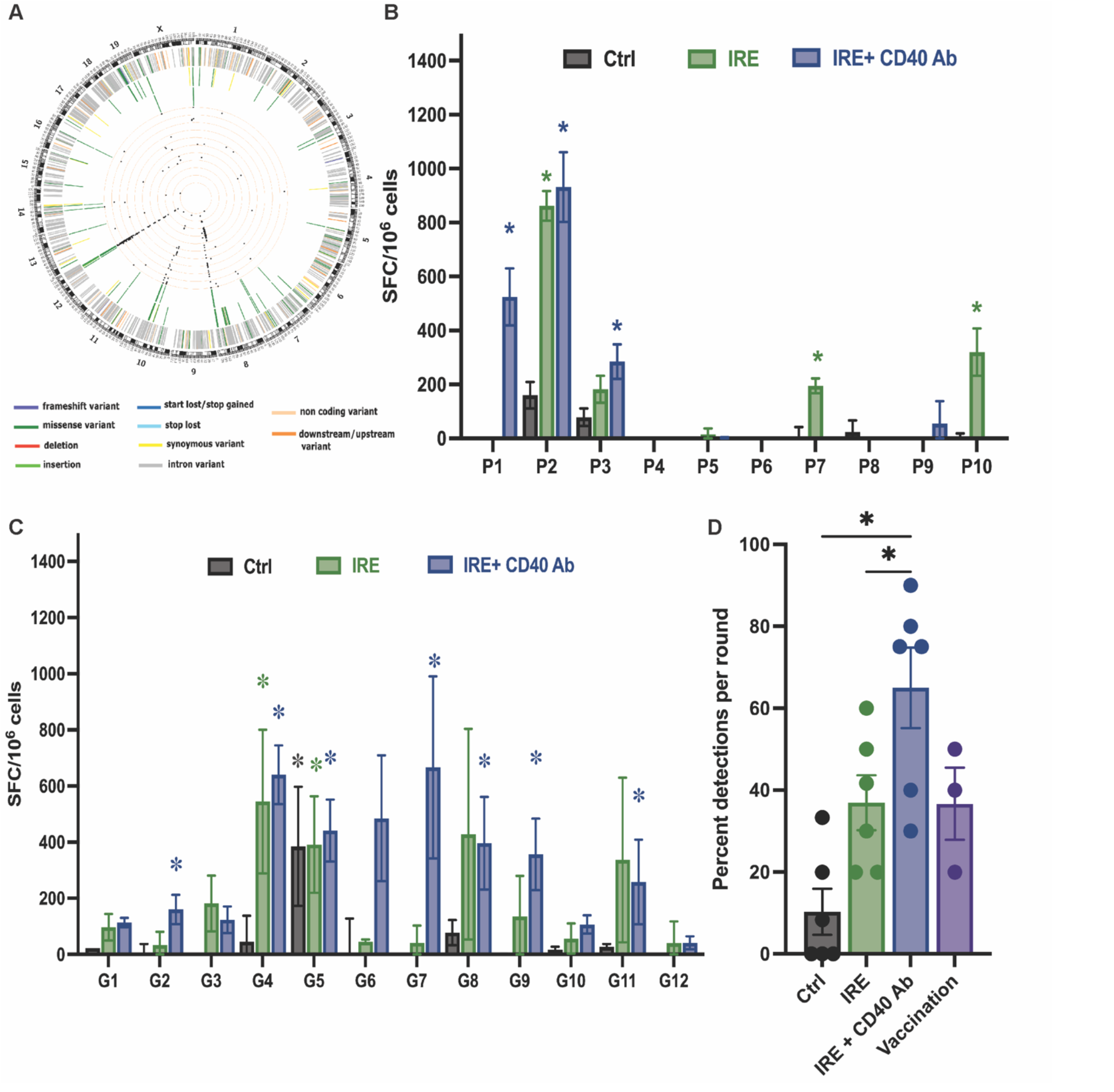
Combination of IRE and CD40 Ab triggers expansion of systemic tumor neo-antigen specific T-cell response. A - Circos plot showing the observed mutations in the KPC46 O tumor compared to B6/129 F1 hybrid background. First level (right next to cytogenic bands): all of the somatic mutations identified by whole exome sequencing. Second level: mutations expressed by RNAseq. Third level: histogram showing the level of RNA expression. Fourth level: 59 mutations selected for peptide synthesis and ELISPOT based on high RNA expression and sequencing depth. Representative IFNo ELISPOT from (B) peptide pools (10 pools of 8-12 peptides each) or (C) deconvoluted individual peptides of pool 2 using T cells isolated from groups of untreated tumor-bearing mice (Ctrl), IRE-treated mice (IRE) and mice treated with a combination of IRE with IT CD40 Ab (IRE+CD40Ab) rechallenged with live tumor cells. Data represent mean □ SEM values of spot forming cells/10^6 cells from 3 independent mice per group in triplicates. Representative graph of three independent experiments. *, P < 0.05; by two-way ANOVA with post hoc Tukey’s multiple comparison test. D – Measure of breadth of neoantigen detection by T-cells represented as percentage of positive detections (2 standard deviations over the background IFN□ response for that mouse) for each treatment group among total available antigens for that round. Each dot represents a single round of detection of multiple pools/peptides mean □ SEM.

### IRE+CD40Ab combination therapy reduces spatial restriction of tumor infiltrating immune cells

Immunohistochemistry and multiplex immunofluorescence analyses enabled us to perform spatial analyses of the immune cells within liver metastases. We observed that effector immune cells like CD8+ T-cells (Fig. 4A) and activated CD80+CD11c+ dendritic cells (Fig. 5A) were restricted to the periphery of the tumor in the sham group. This phenomenon was reversed following treatment with IRE+CD40Ab, where these effector cells were distributed throughout the tumor site. Not only were there more CD8+ T cells in the liver metastases, but also the density of the CD8+ T-cells increased towards the center of the tumor nodule (Fig. 7A). To quantify this phenomenon, we established the parameter of Mean Distance Ratio (MDR) as described in the Methods section. A value of 1 would indicate all cells clustering at the center of the region of interest, whereas a value of 0 indicates all cells restricted to the periphery. The representative heatmap (Ranging from MDR = 0 blue to MDR = 0.92 Red) in Fig. 7B depicts distance from the perimeter for the selected cell type (T-regs here) using pseudo-colors. Not only were there lower number of T-regs following treatment with the combination, but also their distribution was restricted to the periphery in the IRE+CD40Ab group. In contrast, in the sham group, most of the T-regs were concentrated towards the center of the tumor. The MDR from the periphery was significantly lower following treatment for immunosuppressive cell populations such as T-regs (0.76±0.08 vs 0.17±0.07, Fig., 7D) and MDSCs (0.37±0.05 vs 0.14±0.03, Fig. 7D). Similarly, although macrophages were not further characterized as M1 vs M2, their MDR from the perimeter was significantly decreased with treatment (Fig. 7G). The observation was reversed for effector cells and antigen-presenting cells, which were restricted to the periphery in sham-treated tumors but more uniformly distributed throughout the tumor following treatment. The MDR from the periphery was significantly higher following treatment for CD8+ T cells (0.26±0.02 vs 0.5±0.07, Fig. 7C) and CD80+ CD11c + dendritic cells (0.27±0.03 vs 0.47±0.03, Fig. 7E). Although not statistically significant, the observation that the distance ratio between CD40L clusters and CD11c + dendritic cells decreases with IRE+CD40Ab treatment (0.22±0.12 vs 0.06±0.02) is consistent with an active antigen recognition and presentation in the TME.

**Figure 7.**
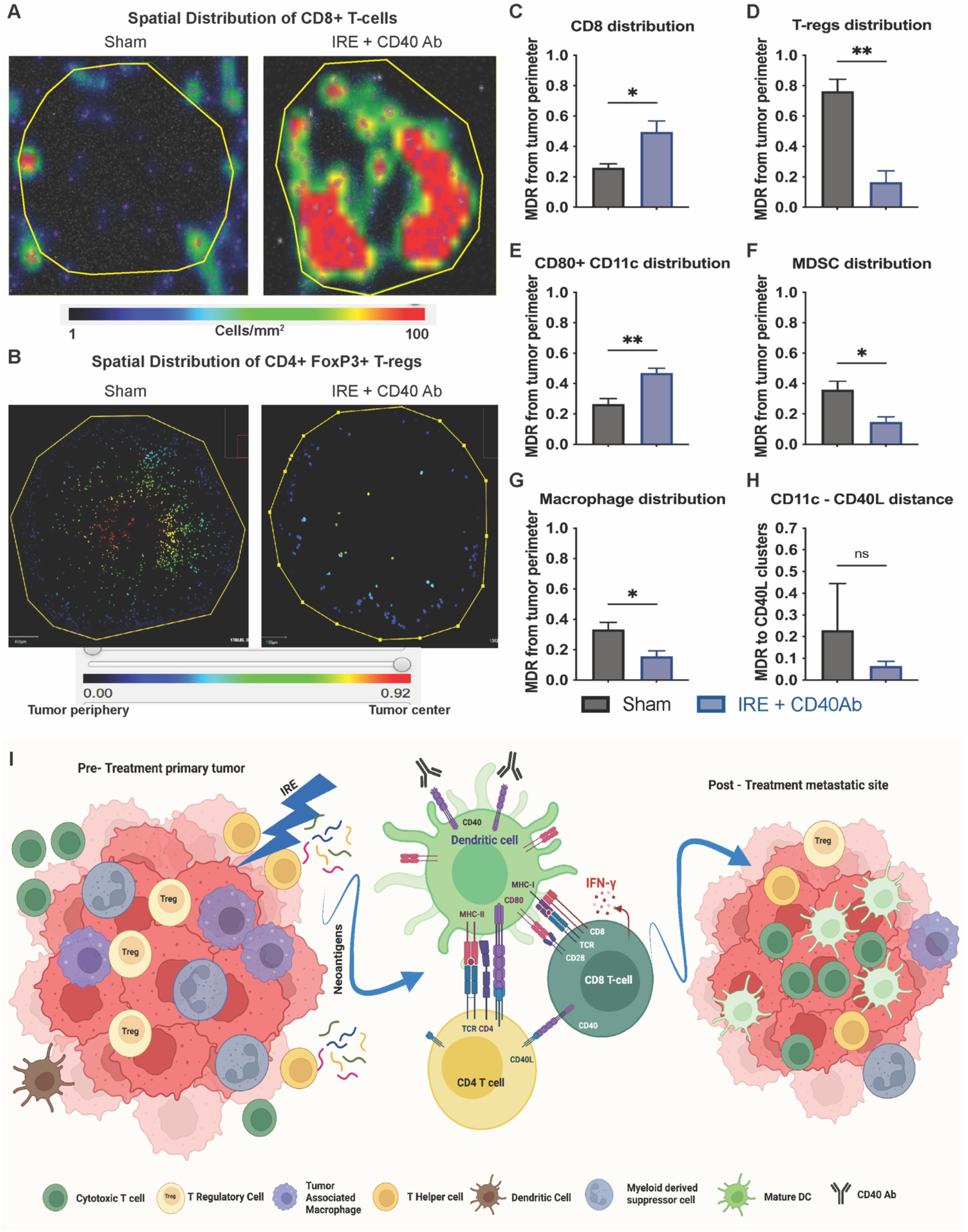
Decreased spatial restriction of infiltrating effector immune cells in the liver observed post treatment with IRE+CD40Ab. A - Representative heat map overlay depicting the density of CD8+ T cells around tumor perimeter shows higher concentration of CD8 T cells within the tumor perimeter post treatment with IRE+CD40Ab. B – Heatmap of spatial analysis of CD4+ FoxP3+ positive regulatory T cell (T – reg) infiltration in the liver metastases with pseudo colors ranging from blue indicating cells closest to the tumor perimeter to red indicating closest to the center of the tumor (Scale MDR = 0 – 0.92). Images show that not only were the number of T - regs reduced upon IRE+CD40Ab treatment but the distribution of the T - regs was restricted to the periphery of the tumor post treatment. (C-G) Spatial distribution of various immune populations were calculated using QuPath. The distance from the defined tumor perimeter was calculated for each cell of interest and normalized to the size of the corresponding node. Mean Distance ratio (MDR) = (o(Distance in om of each cell of interest from tumor perimeter/mean radius of the metastatic node)/Total number of cells of interest). An MDR = 0 represents cells at tumor perimeter and an MDR =1, represents cells at the farthest distance from the perimeter (i.e., the centroid of the tumor). MDR values for at least 3 different biological replicates was calculated and the data presented as mean ± SEM. *, P < 0.05; **, P < 0.01; ***, P < 0.001; by two-tailed student t test of Sham vs IRE+CD40Ab. H – MDR Measures the distance between a CD11c+ cell and a CD40 L expressing cluster as a measure of proximity between the markers, which is needed for DC maturation. I – a schematic representation showing IRE of pancreatic tumors releases tumor specific neo-antigens. Enhanced maturation and activation of dendritic cells by the addition of CD40 Ab enables a broader recognition of these neoantigens thereby resulting the activation of a systemic anti-tumor immune response, this can be evidenced by an increased infiltration of effector immune cells and a restricted infiltration of immunosuppressive cells at the distant metastatic sites in the liver.

## DISCUSSION

Pancreatic cancer (PC) is a systemic disease. Even patients with small, resectable tumors will usually develop distant progression after seemingly complete resection and optimal adjuvant chemotherapy [37] There is intense interest in developing adjuvant immune strategies that can eradicate micrometastatic disease. It is, however, challenging to model micrometastatic disease and test adjuvant therapy strategies in mice. Our orthotopic organoid model consistently develops visible liver metastases between 4-5 weeks after implantation. Before the development of visible metastatic disease, there is a window in which the primary tumor in the pancreas is palpable but “treatable” (< 200 mm^3^) and well under ethical guidelines for humane endpoints. It is therefore a robust tool to study local therapies such as ablation and radiation in combination with adjuvant immunotherapy.

We have previously shown that IRE alone can generate NeoAg-specific T-cells and T-cell-mediated protective immunity in a SQ model of PC [24]. Scheffer et al. demonstrated that IRE can also generate NeoAg-specific T-cell reactivity in human patients [20]. *De novo* T-cell reactivity to the rationally-selected PC NeoAg Wilms Tumor-1 (WT1) was detected in two of seven patients without pre-existing reactivity; moreover, “boosting” of reactivity in two of three patients with pre-existing reactivity also was observed [23]. However, most patients who undergo IRE for locally advanced PC ultimately develop distant metastatic disease [12, 15-17, 38]. Therefore, it is apparent that the immune effects of IRE alone are not sufficient to eradicate distant micrometastatic disease in humans.

Rationale clearly exists for combining IRE with immunotherapy to improve its systemic immune effects, but the optimal agents are not yet clear. Using an orthotopic PC model, Zhao et al. demonstrated that IRE could reverse resistance to anti-PD1 checkpoint inhibition, prolonging survival and promoting infiltration of CD8+ T cells [21]. A Phase II clinical trial of IRE with adjuvant PD-1 checkpoint inhibition (NCT03080974) demonstrated that this combination is well-tolerated and results in increased circulating effector memory T-cells at 90 days [39]. Nevertheless, given the lack of success of checkpoint inhibitors in PC to date, additional strategies—such as stimulation of the innate immune system—are likely to be necessary for effective therapeutic immunity.

CD40 agonists activate dendritic cells and stimulate antigen presentation to T-cells, qualifying them as potentially better single agent adjuvants than checkpoint inhibitors, which require the myeloid arm of the immune system to be engaged separately through the innate immune system. The IRE procedure provides an opportunity to access the tumor, and IT injection delivers CD40Ab directly to the site of antigens released by IRE and the antigen-presenting cells with which they interact. In a bilateral flank SQ model, we have demonstrated that a single IT injection of CD40 Ab at the time of IRE on one flank was more effective at inhibiting growth of contralateral tumors (abscopal effects) than the combination of multiple systemic anti-PD-1 and IT TLR7 agonist injections in an almost identical model (Fig. 1B and [24]). We then utilized our orthotopic model to evaluate this “all local” treatment approach. IRE alone but not CD40Ab alone had significant effects on pancreatic tumor growth. Conversely, CD40 Ab alone but not IRE alone had significant effects on metastatic tumor growth. The combination had at least additive effects, suppressing the growth of both pancreatic tumors and metastatic tumors in the liver, significantly prolonging survival. Analysis of gene expression changes within the primary tumor revealed upregulation of pathways involved in antigen presentation with the combination treatment. The combination of IRE and CD40Ab synergistically promoted the infiltration of active dendritic cell and cytotoxic T-cell infiltration into distant liver metastases.

Several studies have demonstrated that it is not just the number of infiltrating immune cells that is important but their spatial distribution within the tumor and relationships to each other. Several studies in primary PC tissue have documented that effector immune cells are often trapped in the peritumoral space [40, 41]. Our model allows analysis of spatial changes within the metastatic niche following treatment with IRE+CD40Ab. We observed a consistent inverse relationship between the distribution of immunosuppressive MDSCs and T-regs and the effector immune population, including CD8+ T cells and activated dendritic cells. Treatment with IRE+CD40Ab resulted in immunosuppressive cells being more restricted to the periphery and effector cells being more uniformly distributed (Fig. 7). Cognate interaction of CD8+ T-cells with tumor cells is critical to their cytotoxic activity, and a more uniform distribution of CD8+ T cells within tumors correlates with better outcomes [42]. We observed closer interactions of dendritic cellss with CD40L clusters following treatment with CD40Ab indicative of active antigen presentation and cross-presentation (Fig. 7H). Further, infiltration and activation of dendritic cells is critical to the expansion of the T-cell repertoire. To our knowledge, this is the first analysis of spatial differences in the infiltration of immune cells within spontaneous liver metastases, which are not typically resected in humans so not as available for evaluation as primary tumor tissue. We corroborated these findings with a parallel analysis of T-cell reactivity to candidate peptide NeoAgs identified by mutanome anslysis. The combination of IRE+CD40Ab increased the number of peptides recognized by T-cells from treated mice (Fig. 6D). This “NeoAg spreading” may represent an expanded tumor-targeted T-cell repertoire and result in more effective killing of cancer cells.

Our study has several limitations. One is that our model does not completely recapitulate IRE in humans, whose tumors are often heavily pre-treated with chemotherapy and/or radiation therapy. Another is that we cannot exclude the possibility that locally-delivered CD40Ab is absorbed systemically, although this is not necessarily a problem, since systemic (intravenous) CD40Ab delivery has been well-tolerated in clinical trials [36]. Another is that our spatial immune infiltration analysis of the liver was limited to a single timepoint (14 days) and did not capture temporal changes in immune cell infiltrates. Longer time-course experiments will be necessary to assess the durability of these responses. However, this “all local” approach does not preclude the use of additional doses of CD40Ab (either local or systemic) or other rational agents as maintenance therapy. Finally, KPC46 organoids were derived from male KPC mice in immunocompetent B6/129 hybrid mice, which is a first filial generation hybrid between C57BL/6 and 129S1/SvImJ mice. However, since it is an F1 hybrid, spontaneous recombination between the mating parents results in slightly different single nucleotide polymorphism (SNP) profiles among the offspring. The “NeoAgs” identified in our KPC46 model tumor are SNPs relative to a representative B6/129 host, and peptides representing these “NeoAgs” may have different immunogenicity in different B6/129 mice. This resulted in our inability to identify a set of peptides that were highly immunogenic in all mice. This, however, is reflective of human PC, which typically has only a low to moderate mutational burden.

In conclusion, using multiple mouse models of PC, including a model of spontaneous liver metastasis, we have shown (Fig. 7I) that IRE alone can induce local tumor regression and release tumor-specific NeoAgs with beneficial but modest effects on infiltrating immune cells. Addition of locally-delivered CD40Ab at the time of IRE improves the recognition of these NeoAgs by activating dendritic cells, thereby generating a stronger systemic anti-tumor T-cell response and inhibiting metastatic disease progression. These data provide strong rationale for a clinical trial in the setting of locally advanced PC, where patients have a high likelihood of micrometastatic disease. Human PC does not have many prevalent, immunogenic NeoAgs that can be targeted with “off-the-shelf” vaccine approaches, requiring more “personalized” vaccine approaches. “In situ” vaccination with IRE is essentially a form of personalized vaccine that requires no knowledge of the patient’s unique NeoAg profile. A clinical system for IRE (marketed as Nanoknife®, Angiodynamics) has 510(k) clearance from the FDA and is currently being used for selected patients with locally advanced PC who have not developed distant progression after neoadjuvant chemotherapy. A first-in-human study of IT injection of an agonistic CD40 Ab, ADC-1013 or mitazalimab (Alligator Biosciences), has demonstrated that injection even into deep tumors (mostly liver) was feasible and safe [43]. A clinical trial combining IRE with local delivery of CD40 Ab would therefore be imminently feasible. This approach could help improve outcomes for patients with locally advanced PC in the near-term. If the combination of IRE with immunotherapy proves to be effective at decreasing recurrence in patients with locally advanced PC, then this concept could logically be extended to patients with limited metastatic disease.

## MATERIALS AND METHODS

### Cell lines and Organoids

The male KPC4580P cell line was established from a spontaneous tumor that developed in a male LSL-*Kras*^G12D/+^; LSL-*Trp53*^R172H/+^; *Pdx1*^Cre/+^; LSL-*Rosa*26 ^Luc/+^ (KPC-luc) mouse as previously described (gift of Jen-Jen Yeh, UNC [28]). The cells were grown in DMEM:F12 containing 10% fetal bovine serum and 1% antibiotics (Penicillin:Streptomycin) at 37° C with 5% CO_2_. The cell line was authenticated by sequencing and confirmed negative for pathogens using IMPACT testing (IDEXX Bioresearch). KPC-46 organoids were derived from a male *Kras*^*LSL-G12D*^*-p53*^*LSL-R172H*^*-Pdx-1-Cre (KPC)* using previously described methods (gift of Andrew Lowy, UC San Diego [44]). In short, ∼200-300mg of primary tumor tissue was washed in PBS, minced into small pieces and added to 4.7ml RPMI with 1 mg/mL Collagenase and dispase and incubated for 1 h at 37° C. The enzymes were then removed by centrifugation and the cells were placed in 12-well tissue culture dish at a density of 100,000 – 200,000 cells per 50 uL of growth factor-reduced matrigel. 800 uL of growth media containing RPMI, 5% FBS, 2X P+S, 1mM Glutamax, 1mM sodium pyruvate, 1X NEAA, 1X Fungizone, 5ug/ml insulin, 1.4uM hydrocortisone, 10ng/ml EGF, 10.5uM rho kinase inhibitor.

### Animals

All animal experiments were approved by the Institutional Animal Care and Use Committee (IACUC) of University of California, San Diego (UC San Diego). All methods involving animals were performed according to Office of Laboratory Animals Welfare (OLAW) – NIH guidelines, in a facility fully accredited by the Association for Assessment and Accreditation of Laboratory Animal Care, International (AAALAC). 6-8 week old wild type (WT) C57BL/6 and B6/129 F1 hybrid mice were purchased from Jackson Laboratories (Bar Harbor, ME).

### IRE in subcutaneous mouse models

Subcutaneous (SQ) pancreatic tumors were initiated by implanting 5×10^5^ KPC4580P cells in the left flank of 6-8 week old male (gender-matched to cell line of origin) C57BL/6 mice. IRE was performed when tumors reached 4-5 mm diameter, using an ECM 830 square wave pulse electroporator (Harvard Apparatus, Holliston, MA) with a 2-needle array probe, separated by 5 mm, to deliver a total of 150 pulses at 1500 V/cm as previously described [24]. Tumor rechallenge was performed on complete responders with SQ injection of 5×10^5^ KPC4580P cells on the contralateral (right) flank. Age-matched C57BL/6 male mice with a single tumor challenge were used as controls for all rechallenge experiments. Short-term and long-term protective anti-tumor immunity were tested by rechallenge 20 and 200 days post procedure, respectively. SQ tumor sizes were measured using calipers along 2 dimensions and tumor volume (V) was calculated using the formula V = (L x W^2^)/2, where L is the longer and W is the shorter dimension.

### IRE in orthotopic mouse models

For the orthotopic organoid model, solubilized basement membrane matrix (Matrigel, Corning, NY) domes containing the organoids were dislodged from the culture dish and resuspended in 25 mL of cold media. The organoids were then sheared out of the Matrigel scaffold using 23 G needle to establish single organoid suspensions. A small portion of the suspension (1 – 2 mL depending on the extent of organoid growth) was retrieved, and centrifuged at 2000 RPM for 15 min. To the pellet, 1 mL of TryPLExpress cell dissociation reagent was added and was incubated 1 h at 37° C to achieve single cell suspension. A cell count was performed on this suspension to establish the total number of cells in the remaining organoid suspension. The suspension containing the organoids was then centrifuged 2000 RPM for 15 min at 4° C, supernatant was discarded carefully without disrupting the pellet, and the organoids were resuspended in 100% growth factor depleted matrigel at a density of 2.5×10^6^ cells/ mL of Matrigel. 20 μL of organoids suspended in Matrigel were injected into the pancreatic tail of 8 week-old male B6/129 F1J mice via laparotomy as described above.

Orthotopic tumor growth was monitored using ultrasound evaluation (SonoQue L5P handheld ultrasound) until tumors reached 3 - 4 mm in diameter. A second laparotomy was performed to externalize the tumor, and IRE was performed using tweezer-style electrodes (TweezerTrode, BTX). The distance between the electrodes and voltage were adjusted to the dimensions of the tumor to achieve 1500 V/cm, and 150 pulses of electricity were delivered [29]. Intratumoral injections of 20 μL of 2.5 mg/mL agonistic rat anti-mouse CD40Ab (InVivoMAb, clone FGK4.5, BioXCell) were performed immediately after IRE, both in the SQ as well as orthotopic tumor models. Control mice underwent sham laparotomy in all experiments involving orthotopic pancreatic tumors. Mice were administered 1 mg/kg buprenorphine before completion of each laparotomy.

### Bulk RNA sequencing and analysis

Tumors from mice 14 days post-procedure were harvested into Trizol and homogenized immediately post-euthanasia. RNA isolation and purification were performed using RNeasy mini kit (Qiagen, Hilden, Germany) according to manufacturer’s instructions. RNA integrity and quantitation were assessed using the RNA Nano 6000 Assay Kit of the Bioanalyzer 2100 system (Agilent Technologies, CA, USA). Library preparation and sequencing were performed by Novogene Co, Ltd (Sacramento, CA). A total amount of 1 μg RNA per sample was used as input material, and Sequencing libraries were generated using NEBNext® UltraTM RNA Library Prep Kit for Illumina® (NEB, USA). Index codes were added to attribute sequences to each sample. In order to preferentially select cDNA fragments 150∼200 bp in length, the library fragments were purified with AMPure XP system (Beckman Coulter, Beverly, USA). PCR products were purified (AMPure XP system), and library quality was assessed on the Agilent Bioanalyzer 2100 system. The clustering of the index-coded samples was performed on a cBot Cluster Generation System using PE Cluster Kit cBot-HS (Illumina). Paired-end sequencing was performed on an Illumina platform, and paired-end reads were generated. Data were analyzed with a HyperScale architecture (https://rosalind.bio/) developed by ROSALIND, Inc. (San Diego, CA). Quality scores were assessed using FastQC. Reads were aligned to the Mus musculus genome build mm10 using STAR. Individual sample reads were quantified using HTseq and normalized via Relative Log Expression (RLE) using DESeq2 R library. DEseq2 was also used to calculate fold changes and p-values and to perform optional covariate correction [45]. Clustering of genes for the final heatmap of differentially expressed genes was done using the PAM (Partitioning Around Medoids) method using the fpc R library. Hypergeometric distribution was used to analyze the enrichment of pathways, gene ontology, domain structure, and other ontologies. The RNA-seq data were uploaded to “NCBI – SRA database” (Accession number : SUB12118538)

### Analysis of tumor-infiltrating immune cells

Mice bearing SQ or orthotopic tumors were euthanized on day 14 post-procedure and approximately 100 mg of the tissue (tumor or liver) was dissociated into a single cell suspension using 1 mg/mL solution of collagenase/dispase (MilliporeSigma) for 40 min at at 37° C. The cells were then filtered through a 70 μm strainer and viability was assessed using ViCell cell counter (Beckman-Coulter). Single cell suspensions containing 3×10^6^ cells/sample were stained using appropriate fluorescent antibody cocktails (Listed in Supplementary Table S1) after Fc blocking and analyzed using flow cytometry (BD FACS Celesta/Novocyte Advanteon). Cells were fixed and permeabilized using intracellular staining reagents (Intracellular Fixation & Permeabilization Buffer Set, eBioscience) for the staining of FoxP3. Data analysis was performed using Flow Logic software (Inivai Technologies).

### Immunohistochemistry (IHC)

Tissue sections of 5 micron thickness were baked at 60° C for 1 h and were cleared and rehydrated through successive alcohol immersion [Xylene (3 times), 100% EtOH (2 times), 95% EtOH (2 times), 70% EtOH (2 times), then deionized water]. Antigen retrieval was performed in Antigen Unmasking Solution (Citrate Based, pH6, Vector, H-3300) at 95° C for 30 min. Hematoxylin and eosin staining were performed for histological analysis on serial sections of tissues. IHC staining was performed on Intellipath Automated IHC Stainer (Biocare) with the following antibodies: anti-CD45 (Rabbit; AbCAM ab10558; 1:200) and anti – CD8 (Rat, Invitrogen 14-0195-82; 1:100) for 1 h. The slides were washed with 2X Tris-Buffered Saline with 0.1% Tween-20 (TBST) and incubated in secondary antibody, anti-Rat HRP Polymer (Cell IDX, 2AH-100) or anti-Rabbit HRP Polymer (Cell IDX, 2RH-050) for 30 min. The tissues were washed 2X in TBST and developed with DAB (brown) Chromogen (VWR, 95041-478) for 5 min and washed again 2X in deionized water. Brightfield images were obtained using Leica Aperio Slide Scanner using 20X objective and the images were analyzed and quantified using QuPath 3.0 software [46].

### Multiplex immunofluorescence assay

5 micron thick sections of FFPE liver tissues from at least 3 biological replicates from each treatment group were deparaffinized and rehydrated as described in the IHC section. Immunofluorescence staining was performed on Intellipath Automated IHC Stainer (Biocare), with the following steps. Peroxidase block with Bloxall (Vector, SP-6000) for 10 min followed by two washes in TBST and blocking with 3% Donkey Serum for 10 min. Primary antibody (List of IF antibodies, Supplementary Table. S2) incubations were carried out sequentially for 1 hour followed by two washes in TBST and incubation with corresponding secondary antibody, anti-Rat HRP Polymer (Cell IDX, 2AH-100) for 30 min. The tissues were then washed twice in TBST and developed using Tyramide Reagent (Tyramide 488, 555, or 647, ThermoFisher, or Tyramide 790, AAT Bioquest) for 10 min and washed again 2X in deionized H2O. The same steps from antigen retrieval to development with Tyramides was repeated for each of the primary - secondary antibody combinations. Tissues were counterstained with DAPI (1μg/ml) for 15 min and mounted on to coverslip with Vectashield Vibrance/w DAPI (Vector, H-1800-10).

The slides were imaged using Zeiss Axio Scan Z1 slide scanner with a 20x 0.8NA objective, Colibri7 light source, and high-efficiency filter sets. Whole slide images were analyzed using QuPath 3.0 software with individual fluorescence channels (DAPI, 488, 555, 647 and 790) set at constant thresholds across images and groups. Cell detection was performed using DAPI – nucleus channel. Single measurement classifier was used to define positive and negative cells on each channel, and sequential object classification was performed to detect cells positive one or more specific markers. Tumor nodules in the liver were designated as regions of interest (ROI) using PanCK staining and nuclear density changes on the DAPI channel compared to corresponding normal liver which was then correlated to the H&E stains on serial sections. The number of positive detections for each marker was normalized to the tumor area for each ROI. Distance of each positive detection from the perimeter of the ROI was defined as “Distance from Tumor Perimeter” and was normalized to the mean radius of the ROI to establish Mean Distance Ratio (MDR) = (Σ(Distance in μm of each cell of interest from tumor perimeter/mean radius of the metastatic node)/Total number of cells of interest).

### Neoantigen detection assay

Untreated KPC4-6 orthtopic tumors were excised from euthanized B6/129 F1 J mice 21 days after implantation. DNA and RNA were extracted from the tumor tissue using DNeasy mini kit (Qiagen) and RNAeasy mini kit (Qiagen). Whole blood from the tail vein and tail snips from B6/129 F1 J mice were collected for DNA and RNA extraction to be used as reference genome. Whole exome sequencing and mRNA sequencing were performed using miSeq platform (Illumina) by Novogene Technologies, Inc (Sacremento, CA). Expressed non-synonymous genetic variants present in the tumor were identified by cross-referencing the DNA and RNA seq data against B6/129 genome (NCBI SRA accession # SUB12107273). Variants were prioritized according to their ability to be presented by MHC molecules using prediction algorithms [47]. Peptides (20 amino acids in length) harboring these potential antigens at positions 6 and 15 were synthesized by TC peptide Labs (San Diego, CA) and separated into peptide pools of 12 peptides each (List of KPC-46 peptides, Supplementary Information). Bone marrow derived dendritic cells (BMDCs) were generated from age and sex matched B6/129 mice as described earlier [28]. Briefly, tibia were excised asceptically, and the bone marrow was collected. Cells were grown at a concentration of 1 × 10^6^ cells/mL in media containing 20 ng/mL of IL-4 and GM-CSF, for 6 days and BMDCs in suspension were collected on day 7. ELISPOT assay was performed as described earlier [24]. Briefly, The BMDCs were incubated with 5 μg/mL of mutant peptides pools to facilitate antigen presentation for 1 day at 37° C. A pre-wet multiscreen-IP filter plate (Millipore) was coated with IFNγ capture antibodies (AN18; Mabtech). 2 × 10^5^ lymphocytes isolated from treatment-responsive mice were incubated with 5 μg/mL of the different mutant peptides pools to which 20,000 activated BMDCs were added. 5 μg/mL of concanavalin A (Sigma-Aldrich) was used as the positive stimulus control and no peptide wells were used as negative control. Lymphocytes from mice subcutaneously injected with irradiated 5×10^6^ KPC-46 cells followed by live cell rechallenge were used as vaccination controls (Vaccinated). IFNγ secretion was assessed biotinylated anti-mouse IFNγ (R4-6A2; Mabtech) and imaged using an ELISPOT reader (AID Diagnostika). Wells with >2 standard deviations more IFNγ reactive spots than the negative control were considered positive.

### Graphics

The graphical representations, schematics and timelines used in the figures were created using BioRender.com.

### Statistical Analysis

All results were expressed as means ± standard error of the mean (SEM). Statistical difference between groups was calculated either using the student’s *t* test or ANOVA with post-hoc multiple comparisons depending on the data, using GraphPad Prism 9.0 software. A value of P < 0.05 was considered significant.

## Supporting information

List of KPC-46 peptides

## Data Availability

All data relevant to the study are included in the article, supplementary data or uploaded to NCBI SRA database. Other raw data are available upon reasonable request.

## CONTRIBUTIONS STATEMENT

JSSN performed the experiments. JSSN and RRW designed the experiments. JSSN, TH, AM, DC, SPS and RRW analyzed and interpreted the data. TH, SE, SMA, HT, PR and ZM provided technical and material support for the experiments. MP and KM provided support for statistical analysis. JSSN and RRW wrote the manuscript. TH, ZM, PR, DC and SPS edited the manuscript. All authors reviewed the manuscript. RRW conceived and supervised the study.

## ACKNOWLEDGEMENTS

Histology support was provided by Biorepository and Tissue Technology Shared Resource (BTTSR) of Moores Cancer Center, UCSD. Bioinformatics support for mutanome analysis was provided by Ashmitaa Premlal and Jason Greenbaum of Bioinformatics core facility at La Jolla Institute of Immunology.

## FUNDING SUPPORT

Research presented in this manuscript was supported by NIH R01CA254268 and the San Diego Cancer Centers Council Padres Pedal the Cause (#PTC2017). The UCSD Moores Cancer Center Biorepository and Tissue Technology Shared Resource is funded by the NIH (P30CA23100). The content is solely the responsibility of the authors and does not necessarily represent the official views of the NIH.

## Notes

### Competing Interest Statement

The authors have declared no competing interest.

