## Supplementary Fig. 1 for "Treatment of pancreatic cancer with irreversible electroporation and intratumoral CD40 antibody stimulates systemic immune responses that inhibit liver metastasis in an orthotopic model"

**Supplementary Data**

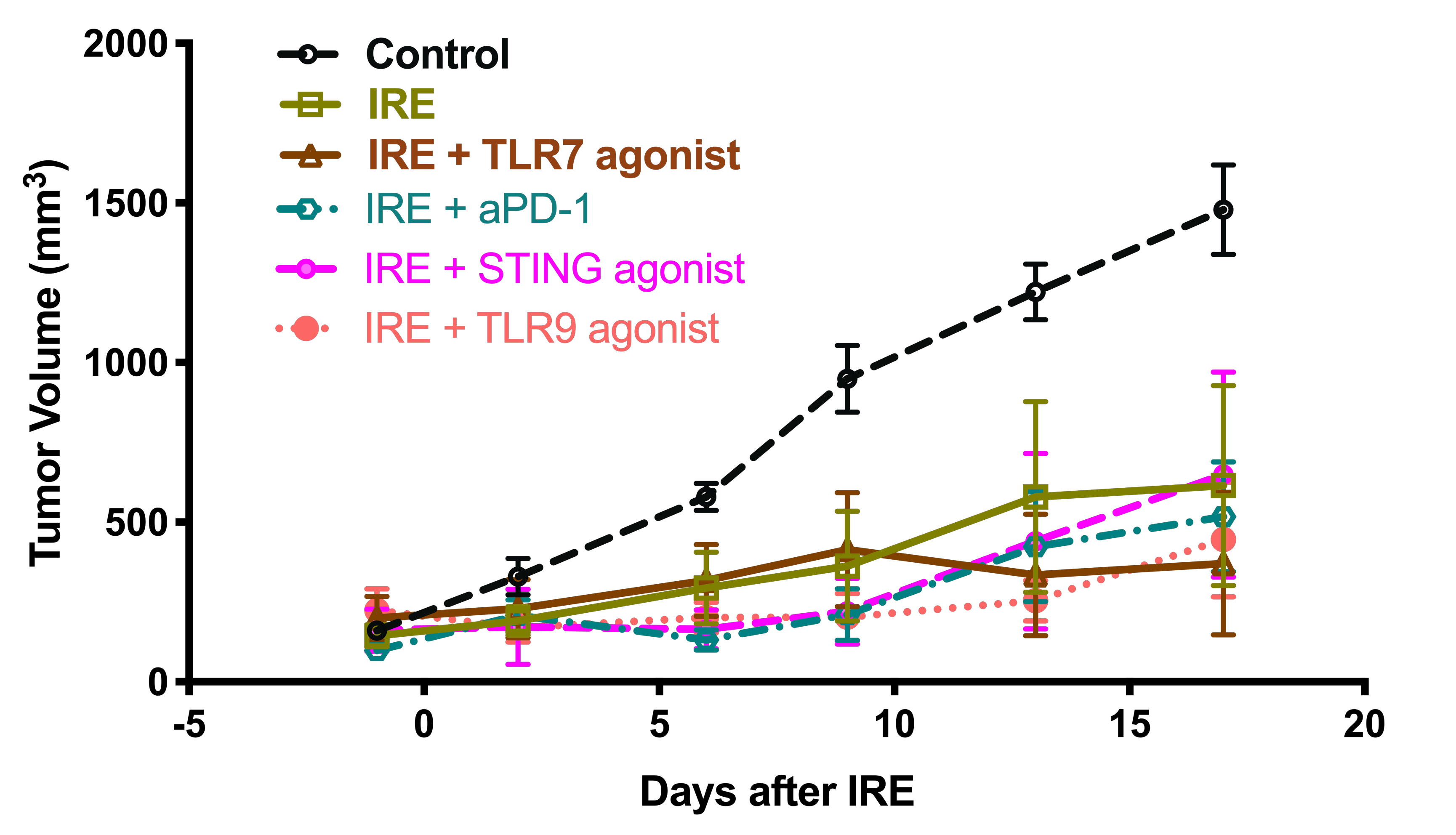

**Supplementary Figure 1. Subcutaneous KPC4580P tumor growth following IRE + various immunomodulatory agents.**

KPC4580P tumor cells (5 × 10^5^ cells/mouse) were subcutaneously implanted on the flanks of C57BL/6 mice 8 days before IRE. IRE was performed on day 0. Intra-tumoral TLR7- or TLR9- or STING agonist or intraperitoneal (IP) anti-PD-1 antibody was given on Days 0, 3, 5 and 7. Tumor volumes were measured using calipers and plotted as mean ± SEM (n=5 mice/group). No significant difference between IRE alone or any of the combination groups was observed.

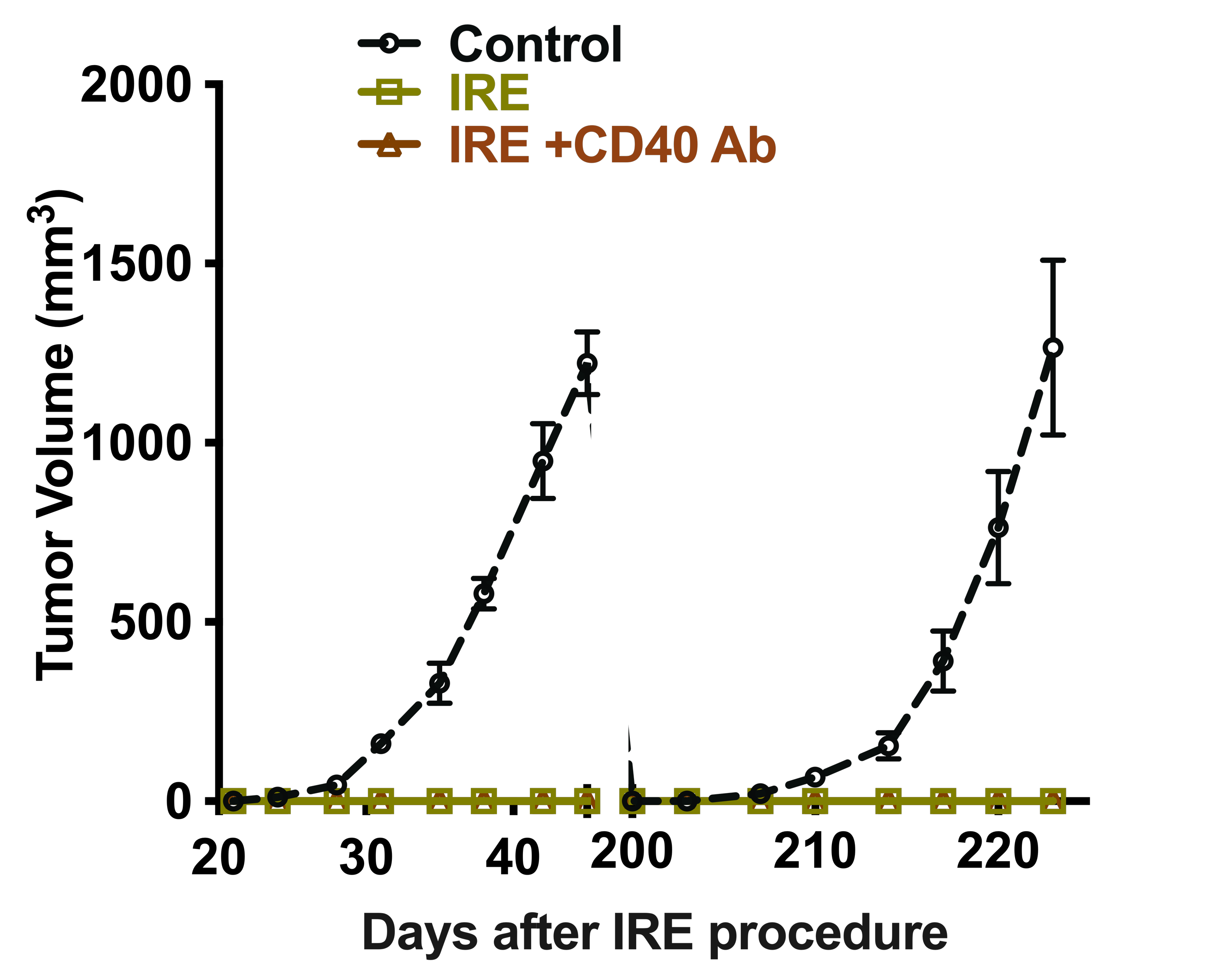

**Supplementary Figure 2. Rechallenge tumor growth in SQ KPC4580P model.**

The complete responders to IRE alone (n=3) or IRE+CD40Ab (n=4) treatment were implanted with tumor cells (500,000/mouse) at 20 (early) and 200 (delayed) days post IRE. Tumor growth was observed for up to 30 days. Tumor growth in age-matched naive mice served as a positive control. Each data point represents mean tumor volumes ± SEM.

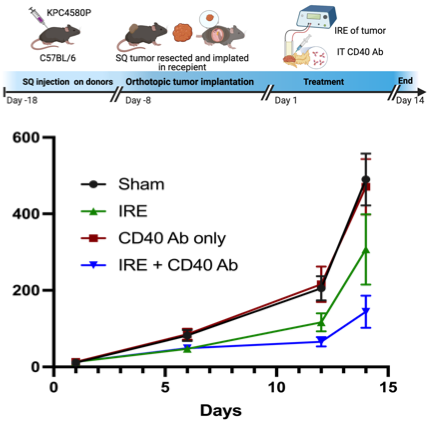

**Tumor Volume (mm3)**

**Supplementary Figure 3. Tumor growth curves of orthotopically-implanted KPC4580P pancreatic tumors**.

C57BL/6 mice were implanted with 1 mm^3^ tumor pieces from SQ KPC4580P donor tumors onto the pancreas. Treatment was performed on day 8 post-implantation. Tumor volumes were measured using ultrasound and plotted as mean ± SEM (n=5 mice/group).

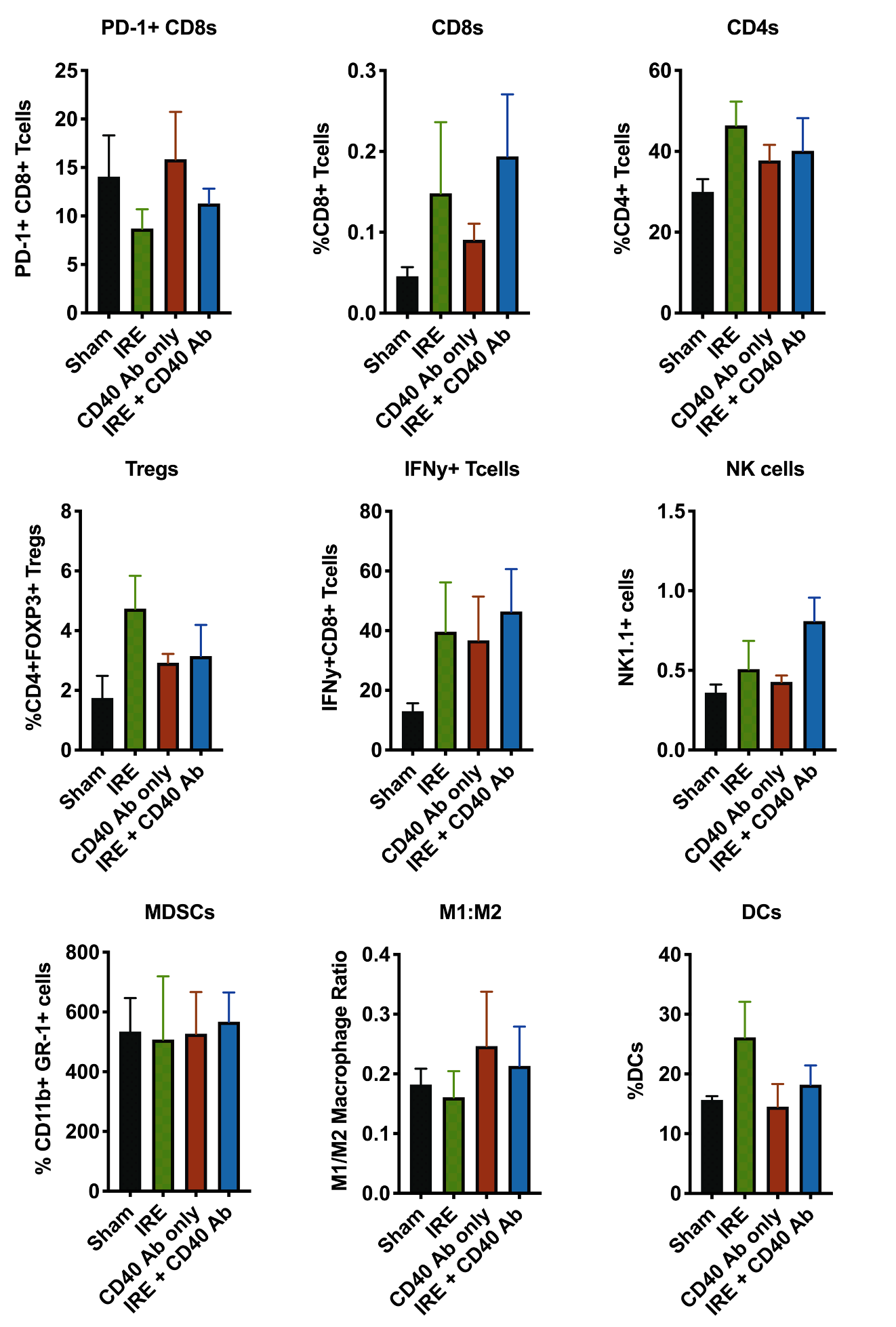

**Supplementary Figure 4. Flow cytometry analysis of primary orthotopic KPC-46 organoid pancreatic tumors**.

Orthotopic KPC-46 organoid pancreatic tumors (n=4 mice/group) were dissected on day 14 post-treatment with IRE+CD40Ab. The single-cell suspension was prepared and stained with appropriate antibodies. The gating strategy and the reagents are shown in Supplementary Figure 7, Supplementary Table 1, and Material and Methods section.

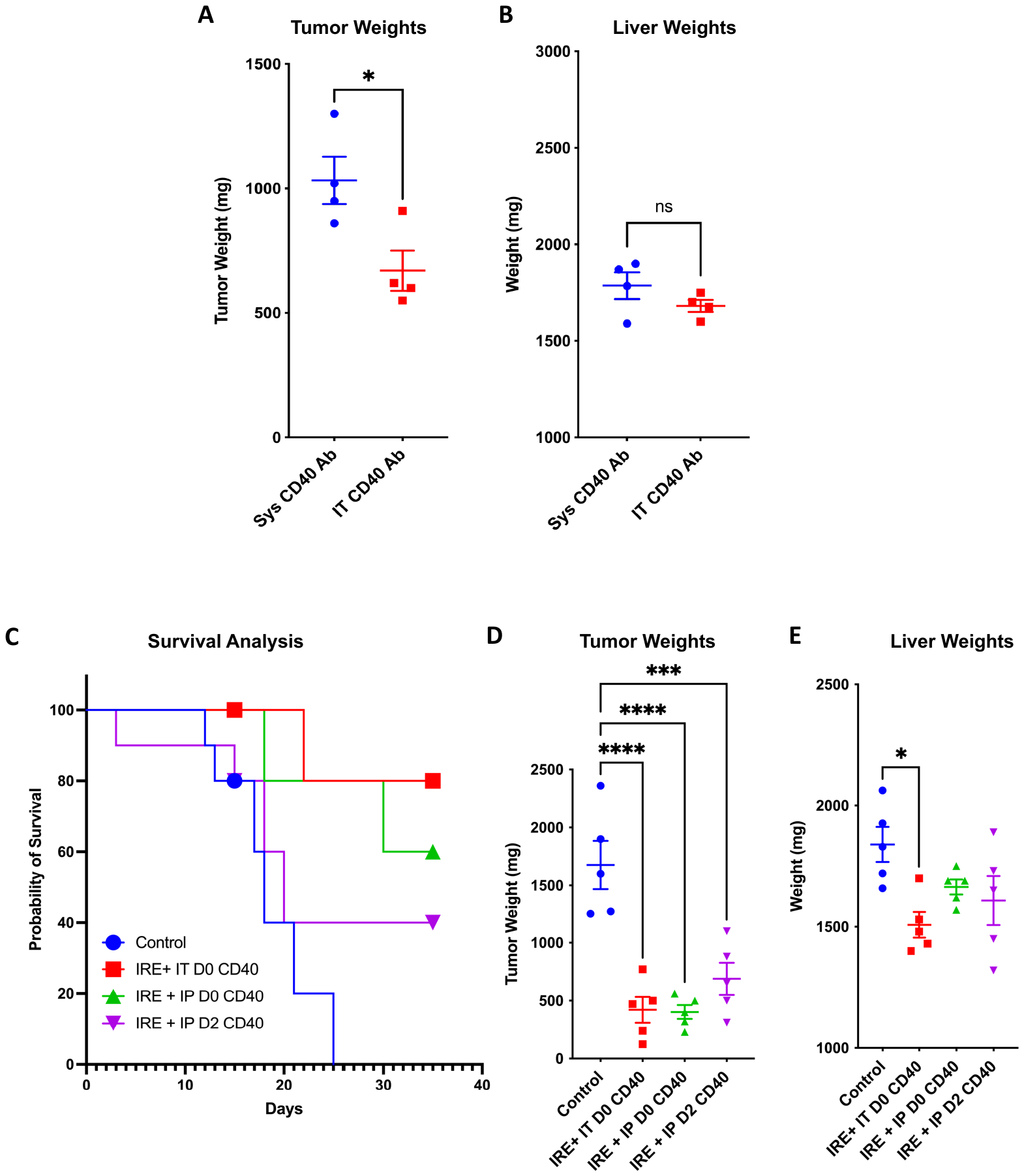

**Supplementary Figure 5. Comparison of delivery routes for CD40 Ab.**

KPC-46 organoid orthotopic pancreatic tumors (n=4/group) were treated with systemic (Intraperitoneal) CD40Ab on days 0 or 2 or intratumoral CD40Ab treatment as a single agent. Tumor-bearing untreated mice served as controls. (A) Primary tumor weights were compared between systemic (IP) and IT routes on day 14 post-treatment. Tumor weight is represented as mean ± SEM. IP injection of CD40Ab alone was less effective than IT injection. (B) Liver weights were measured as an indicator of metastatic burden and were not significantly (NS) different. * P < 0.05 by two-tailed T-test.

Further comparisons of effect of delivery route of CD40Ab was performed in combination with IRE. (C-E) Survival Analysis (C), primary tumor weight (D) and liver weights (D) were compared among 4 treatment groups, day of IRE (D0) IT delivery, D0 IP delivery, and 2 days after IRE (D2) IP delivery, control. There was no significant difference between the three treatment groups with respect to primary tumor control, but significant effects on liver weights were only seen in the D0 IT delivery group. *P<0.05, ***P<0.001, and ****P<0.0001, by one-way ANOVA with post-hoc Benferroni testing.

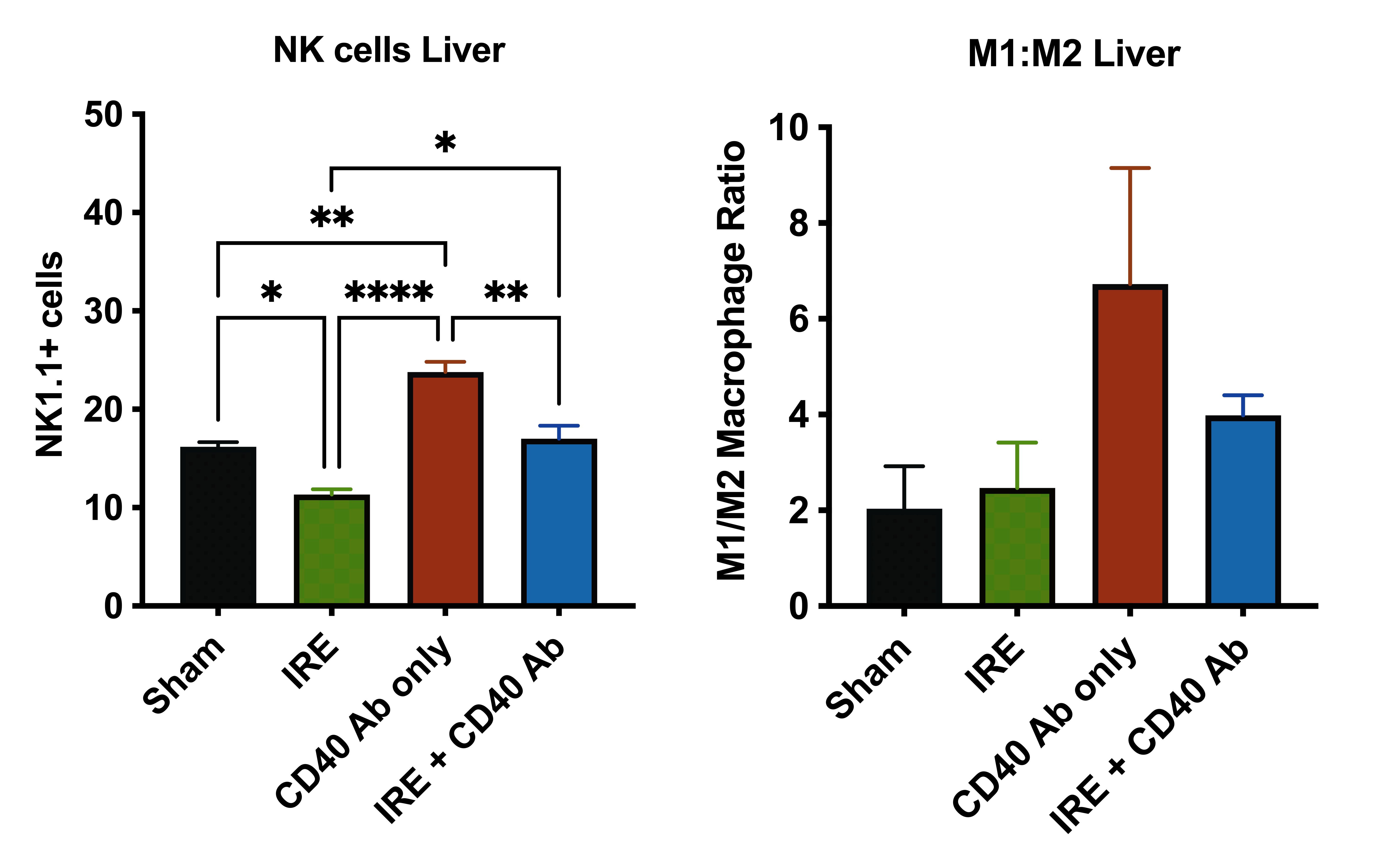

**Supplementary Figure 6. Immune cell infiltration in livers with metastasis.**

Single-cell suspensions of the bulk liver with metastasis (n=3/group) were prepared on day 14 post-treatment. Cells were stained with M1 and M2 macrophage markers (MHC-II and CD206 respectively) and NK cell markers and analyzed by flow cytometry. A trend toward increased M1/M2 ratio and increased infiltration of NK cells suggest anti-tumor immune activity at the distant metastatic site following treatment of the primary pancreatic tumor with CD40Ab. Graphs plotted as mean ± SEM. * P<0.05, *** P<0.001, and **** P<0.0001 by one-way ANOVA with post-hoc Benferroni testing.

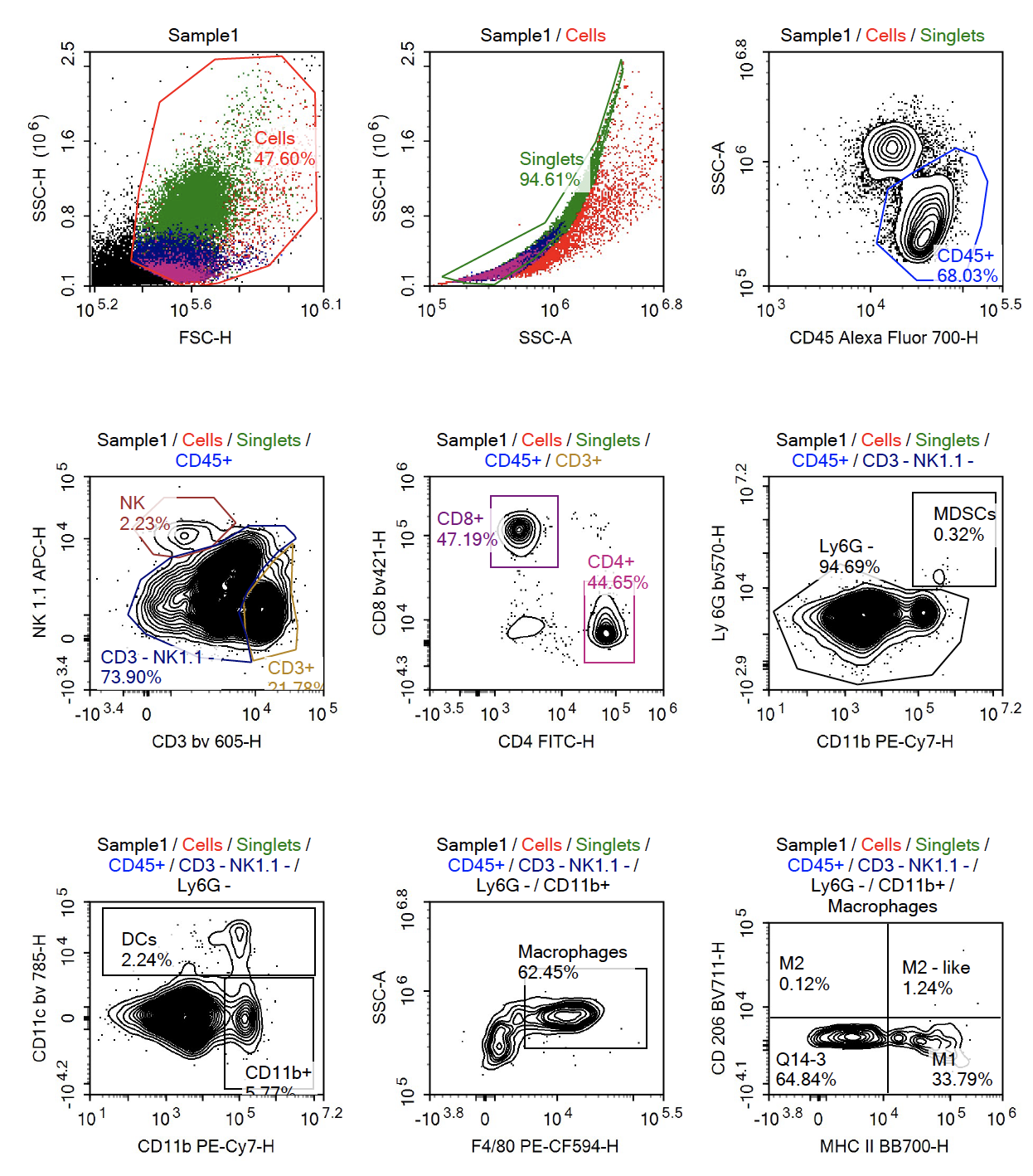

**Supplementary Figure 7. Gating strategy of flow cytometric analysis.**

**Supplementary Table. S1 – List of flow cytometry antibodies**

| **Antibody Target** | **Fluorophore** | **Catalog Number** | **Clone** | **Manufacturer** |
| --- | --- | --- | --- | --- |
| CD11b | FITC | 101206 | M1/70 | Biolegend |
| CD11b | PE-CY7 | 552850 | M1/70 | BD |
| CD11c | APC-CY7 | 561241 | HL3 | BD |
| CD19 | APC-H7 | 560143 | 1D3 | BD |
| CD206 | BV711 | 141727 | C068C2 | Biolegend |
| CD25 | BV650 | 102038 | PC61 | Biolegend |
| CD4 | PE-CY7 | 552775 | RM4.5 | BD |
| CD4 | FITC | 100510 | RM4.5 | Biolegend |
| CD45 | AF700 | 56-0451-82 | 30-F11 | Invitrogen |
| CD8 | PERCP-CY5.5 | 100734 | 53-6.7 | Biolegend |
| CD8 | BV421 | 563898 | 53-6.7 | BD |
| F4/80 | PE-CF594 | 565613 | T45-2342 | BD |
| FOXP3 | AF488 | 320012 | 150D | Biolegend |
| FOXP3 | PE | 560408 | MF23 | BD |
| GR-1 | BV421 | 108434 | RB6-8C5 | Biolegend |
| IFN gamma | BV711 | 564336 | XMG1.2 | BD |
| LY6C | BV650 | 128049 | HK1.4 | Biolegend |
| LY6G | BV570 | 127629 | 1A8 | Biolegend |
| MHC-II | BB700 | 746086 | 2G9 | BD |
| MHC-II | FITC | 35-5321-U500 | M5/114.15.2 | Tonbo |
| NK1.1 | APC | 550627 | PK136 | BD |
| NK1.1 | BV605 | 108753 | PK136 | Biolegend |
| TCR beta | BV605 | 562840 | H57-597 | BD |

**Supplementary Table. S2 – List of Immunofluorescence cytometry antibodies**

| **Antibody Target** | **Source Animal** | **Catalog Number** | **Dilution** | **Manufacturer** |
| --- | --- | --- | --- | --- |
| CD80 | Rabbit | Ab254579 | 1:100 | Abcam |
| Pan-CK | Rabbit | Z0622 | 1:500 | Dako |
| CD11c | Rabbit | 97585s | 1:100 | Cell Signaling |
| CD40 L | Rabbit | Ab65854 | 1:250 | Abcam |
| FoxP3 | Rat | 14-5773-82 | 1:200 | Invitrogen |
| F4/80 | Rat | MCA497B | 1:50 | Bio-Rad |
| Ly6G | Rabbit | 87048s | 1:100 | Cell Signaling |
| CD4 | Rabbit | Ab183685 | 1:100 | Abcam |
